# Homeostatic regulation of REM sleep by the preoptic area of the hypothalamus

**DOI:** 10.1101/2023.08.22.554341

**Authors:** John Maurer, Alex Lin, Xi Jin, Jiso Hong, Nicholas Sathi, Romain Cardis, Alejandro Osorio-Forero, Anita Lüthi, Franz Weber, Shinjae Chung

## Abstract

Rapid-eye-movement sleep (REMs) is characterized by activated electroencephalogram (EEG) and muscle atonia, accompanied by vivid dreams. REMs is homeostatically regulated, ensuring that any loss of REMs is compensated by a subsequent increase in its amount. However, the neural mechanisms underlying the homeostatic control of REMs are largely unknown. Here, we show that GABAergic neurons in the preoptic area of the hypothalamus projecting to the tuberomammillary nucleus (POA^GAD2^→TMN neurons) are crucial for the homeostatic regulation of REMs. POA^GAD2^→TMN neurons are most active during REMs, and inhibiting them specifically decreases REMs. REMs restriction leads to an increased number and amplitude of calcium transients in POA^GAD2^→TMN neurons, reflecting the accumulation of REMs pressure. Inhibiting POA^GAD2^→TMN neurons during REMs restriction blocked the subsequent rebound of REMs. Our findings reveal a hypothalamic circuit whose activity mirrors the buildup of homeostatic REMs pressure during restriction and that is required for the ensuing rebound in REMs.

## INTRODUCTION

REMs is homeostatically regulated as demonstrated in various species including mice, rats, cats and humans (Siegel and Gordon, 1965; Beersma et al., 1990; Benington et al., 1994; Endo et al., 1997, 1998; Rechtschaffen et al., 1999; Franken, 2002; Shea et al., 2008). REMs restriction leads to an increased homeostatic need for REMs. During the subsequent recovery sleep, the amount of REMs is increased to compensate for the amount lost during restriction. While seminal dissection studies indicate a pontine origin of REMs, recent studies have revealed that neural populations in the hypothalamus, midbrain, amygdala and medulla regulate REMs by promoting or inhibiting REMs (Clément et al., 2011; Jego et al., 2013; Hayashi et al., 2015; Van Dort et al., 2015; Weber et al., 2015, 2018; Chung et al., 2017; Torontali et al., 2019). However, we do not have a clear understanding about the homeostatic mechanisms regulating REMs and which brain regions integrate homeostatic REMs pressure.

The preoptic area of the hypothalamus (POA) is crucial for sleep regulation. The POA contains neurons that become activated during sleep and that are sufficient and necessary for sleep (Von Economo, 1930; Nauta, 1946; McGinty and Sterman, 1968; Sallanon et al., 1989; Sherin et al., 1996; Szymusiak et al., 1998; Lu et al., 2000; Gong et al., 2004; Takahashi et al., 2009; Zhang et al., 2015; Kroeger et al., 2018). Specifically, POA GABAergic neurons projecting to the tuberomammillary nucleus (POA^GAD2^→TMN neurons) form a subpopulation of sleep-active POA neurons, and promote sleep when optogenetically activated (Chung et al., 2017). A previous study demonstrated that c-Fos expression in the POA is increased in REMs restricted rats (Gvilia et al., 2006). However, the molecular identity of POA neurons that become activated during heightened REMs pressure and whether their activity is necessary for homeostatic REMs regulation remains largely unknown.

In this study, using fiber photometry, we found that POA^GAD2^→TMN neurons become gradually activated during non-rapid eye movement sleep (NREMs) before the onset of REMs and are most active during REMs. Optogenetic inhibition of POA^GAD2^→TMN neurons significantly decreased REMs. We therefore hypothesized that the POA^GAD2^→TMN neurons are well suited to encode homeostatic pressure for REMs. Using fiber photometry recordings combined with REMs restriction, we show that the activity of POA^GAD2^→TMN neurons significantly increased during periods of heightened REMs pressure. Optogenetic inhibition of POA^GAD2^→TMN neurons during REMs restriction prevented the subsequent increase of REMs during recovery sleep. Our findings identify a hypothalamic circuit regulating the homeostatic need for REMs.

## RESULTS

### POA^GAD2^→TMN neurons are most active during REMs

To monitor the population activity of POA^GAD2^→TMN neurons *in vivo* during spontaneous sleep, we performed fiber photometry recordings. GAD2-Cre mice were injected with retrograde adeno-associated viruses (AAVs) encoding Cre-inducible GCaMP8s (AAVretro-FLEX-jGCaMP8s) into the TMN (Tervo et al., 2016; Zhang et al., 2023), and an optic fiber was implanted into the POA (**Fig. 1A**; virus expression and optic fiber tracts were located in the ventrolateral POA, lateral POA, and the lateral part of medial POA). The calcium activity of POA^GAD2^→TMN neurons was significantly higher during REMs compared with that during wake and NREMs (**Fig. 1B, C**; detailed statistical results are shown in **Figure legends** and **Table S1**).

**FIGURE 1.**
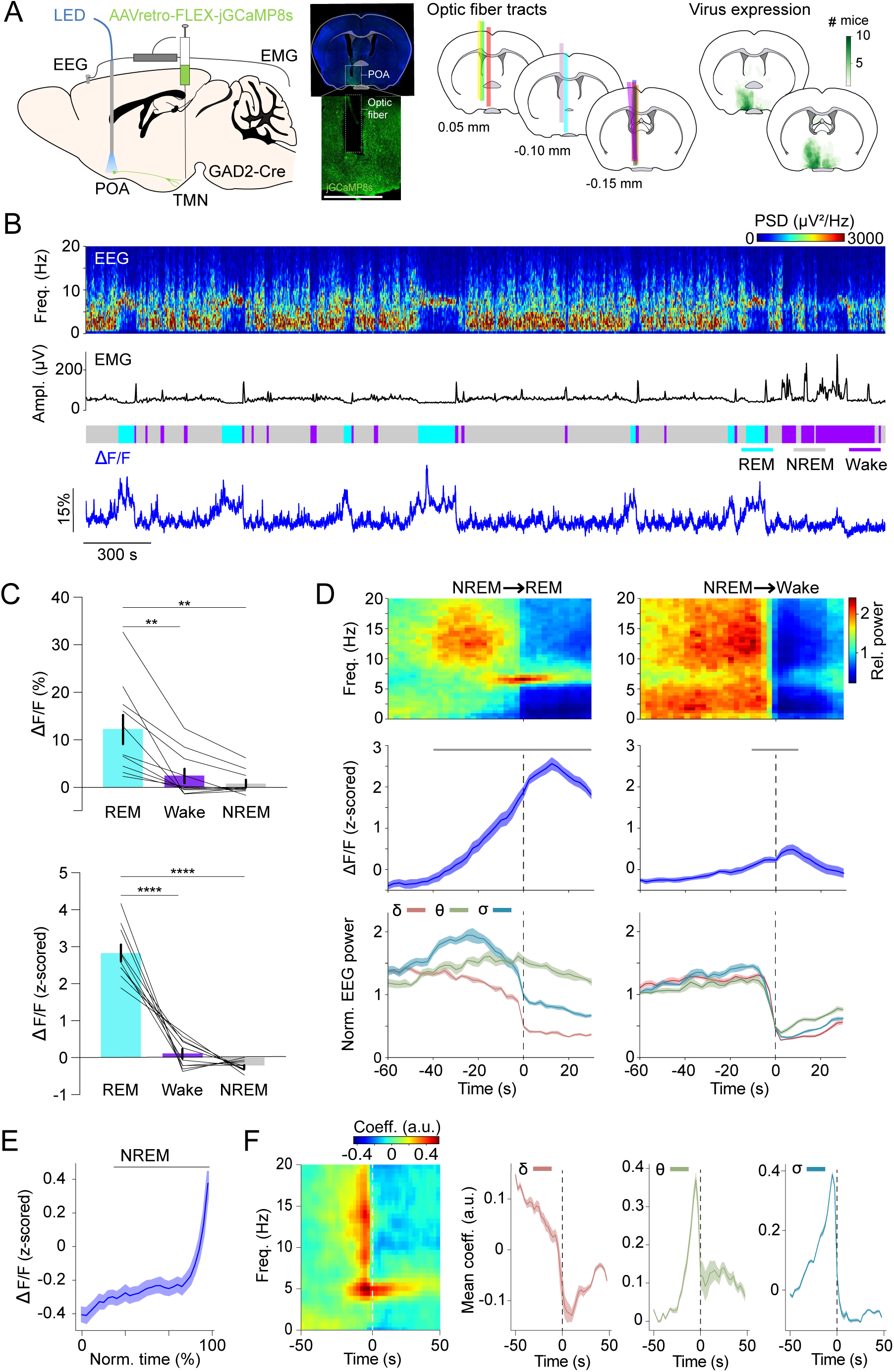
POA^GAD2^→TMN neurons are most active during REMs. **(A)** Left, schematic of fiber photometry with simultaneous EEG and EMG recordings. Mouse brain figure adapted from the Allen Reference Atlas - Mouse Brain (atlas.brain-map.org). Center left, fluorescence image of POA in a GAD2-Cre mouse injected with AAVretro-FLEX-jGCaMP8s into the TMN. Scale bar, 1 mm. Center right, location of fiber tracts. Each colored bar represents the location of optic fibers for photometry recordings. Right, heatmaps outlining areas with cell bodies expressing GCaMP8. The green color code depicts how many mice the virus expression overlapped at the corresponding location (n = 10 mice). **(B)** Example fiber photometry recording. Shown are EEG spectrogram, EMG amplitude, color-coded brain states, and ΔF/F signal. **(C)** Non-normalized and z-scored ΔF/F activity during REMs, wake, and NREMs. Bars, averages across mice; lines, individual mice; error bars, ± s.e.m. One-way repeated measures (rm) ANOVA, p = 9e-4, 2e-6 for non-normalized ΔF/F and z-scored ΔF/F signals; pairwise t-tests with Bonferroni correction, non-normalized ΔF/F, p = 0.0056, 0.0039 for REMs vs. Wake and REMs vs. NREMs; z-scored ΔF/F, p = 6e-5, 1e-7. n = 10 mice. **(D)** Average EEG spectrogram (top), z-scored ΔF/F activity (middle) and normalized EEG δ, θ and σ power (bottom) during NREMs→REMs transitions (left) and NREMs→Wake transitions (right). Shading, ± s.e.m. One-way rm ANOVA, p = 3.18e-49, 9.50e-8 for NREMs→REMs and NREMs→Wake; pairwise t-tests with Holm-Bonferroni correction, NREMs→REMs p < 0.0419 between -40 s and 30 s, NREMs→Wake p < 0.0106 between -10 s and 10 s. Gray bar, period when ΔF/F activity was significantly different from baseline (-60 to -50 s). n = 10 mice. **(E)** ΔF/F activity during NREMs. The duration of NREMs episodes was normalized in time, ranging from 0 to 100%. Shading, ± s.e.m. Pairwise t-tests with Holm-Bonferroni correction p < 8.14e-9 between 20 and 100. Gray bar, intervals where ΔF/F activity was significantly different from baseline (0 to 20%, the first time bin). n = 566 events. **(F)** Left, linear filter mapping the normalized EEG spectrogram onto the POA^GAD2^→TMN neural activity. Time point 0 s corresponds to the predicted neural activity. Right, coefficients of the linear filter for δ, θ and σ power band. Shading, ± s.e.m. See **Table S1** for the actual p values.

We further analyzed the activity changes of the POA^GAD2^→TMN neurons during NREMs→REMs or NREMs→wake transitions. We found that the calcium activity of POA^GAD2^→TMN neurons during NREMs→ REMs transitions becomes significantly increased 40 s before the REMs onset and remains elevated throughout REMs (**Fig. 1D**). During NREMs→wake transitions, the activity of POA^GAD2^→TMN neurons started rising 10 s before the wake onset and remained elevated for 10 s after the onset (**Fig. 1D**). The ΔF/F activity gradually increased throughout NREMs episodes (**Fig. 1E**).

Given that both the θ and σ (6-9 and 10.5-16 Hz) power increased preceding the REMs onset (**Fig. 1D**), we investigated the relationship between the POA^GAD2^→TMN neuron activity and the spectral composition of the EEG in more detail (**Fig. 1F**). We used a linear regression model to predict the current POA^GAD2^→TMN neural activity based on the preceding and following spectral EEG features (Weber et al., 2010; Schott et al., 2023). We found that the activity of POA^GAD2^→TMN neurons is preceded by an increase in the θ and σ power and reduction in the δ (0.5-4.5 Hz) power (**Fig. 1F**). These changes in the EEG are characteristic for the stage of NREMs preceding REMs (Gottesmann, 1996) and their correlation with the activity of POA^GAD2^→TMN neurons is consistent with a role of these neurons in promoting NREMs to REMs transitions.

In a complementary experiment, we monitored the calcium activity of POA^GAD2^→TMN axonal fibers. GAD2-Cre mice were injected with AAV-FLEX-GCaMP6s into the POA and an optic fiber was implanted into the TMN (**Fig. S1A**). Similar to the activity found for POA^GAD2^→TMN neurons using retrograde AAVs (**Fig. 1B, C**), POA^GAD2^→TMN fibers were most active during REMs (**Fig. S1B, C**). During NREMs, the activity of POA^GAD2^→TMN axonal fibers gradually increased before transitioning to REMs (**Fig. S1D**), demonstrating that the activity of POA^GAD2^→TMN axonal fibers closely resembles that of cell bodies. In summary, these results demonstrate that the activity in POA^GAD2^→TMN neurons increases prior to the onset of REMs episodes, following an increase in the θ and σ power in the EEG.

The TMN contains histamine neurons (TMN^HIS^), and previous studies showed that POA^GAD2^ neurons innervate TMN^HIS^ neurons (Chung et al., 2017; Saito et al., 2018). Consistent with previous electrophysiological studies (Steininger et al., 1999; Vanni-Mercier et al., 2003; John et al., 2004; Takahashi et al., 2006), we found in photometry recordings that TMN^HIS^ neurons are highly active during wake and less active during NREMs and REMs (**Fig. S2A-C**). Consistent with this, the activity of TMN^HIS^ neurons became significantly activated after the transition from NREMs or REMs to wakefulness (**Fig. S2D**). The TMN^HIS^ neuron activity gradually decreased during NREMs episodes (**Fig. S2E**), possibly as a result of inhibitory inputs from POA^GAD2^→TMN neurons (Chung et al., 2017). Moreover, examining the time course of TMN^HIS^ neurons between two successive REMs episodes (inter-REM interval), we found that their activity gradually decreases throughout the inter-REM interval, reaching its lowest level at the onset of REMs (**Fig. S2F**). This finding suggests that a minimal activity of TMN^HIS^ neurons is required for entering REMs, and suppression of TMN^HIS^ activity therefore likely facilitates transitions to REMs.

### Inhibiting POA^GAD2^→TMN neurons reduces REMs

To examine whether POA^GAD2^→TMN neurons regulate REMs, we optogenetically inhibited these neurons using the bistable chloride channel SwiChR++ (Berndt et al., 2016; Smith et al., 2022; Stucynski et al., 2022). GAD2-Cre mice were bilaterally injected with retrograde AAVs encoding Cre-inducible SwiChR++ (AAVretro-DIO-SwiChR++-eYFP) or eYFP (AAVretro-DIO-eYFP) into the TMN followed by bilateral optic fiber implantation into the POA (**Fig. 2A**). We compared SwiChR++ and eYFP recordings with and without laser stimulation (473 nm, 2 s step pulses at 60 s intervals for 3 h, zeitgeber time [ZT] 2-5). SwiChR++-mediated inhibition of POA^GAD2^→TMN neurons reduced the amount of REMs compared with recordings without laser stimulation in the same mice and eYFP mice with laser stimulation (**Fig. 2B, C**). The overall time spent in wake and NREMs was not altered by SwiChR++-mediated inhibition (**Fig. 2C, S3A, B**) suggesting that POA^GAD2^→TMN neuron activity specifically regulates the amount of REMs.

**FIGURE 2.**
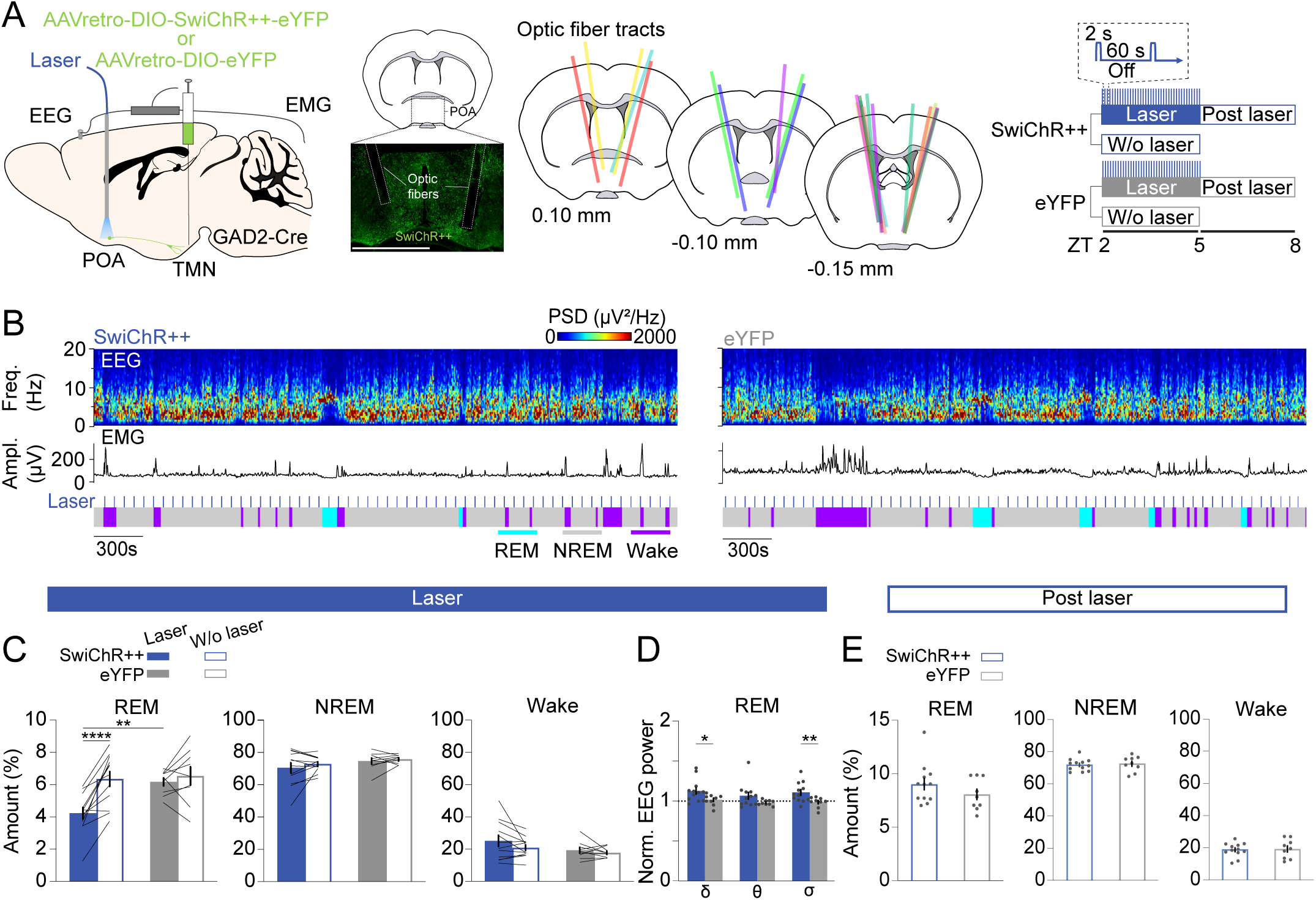
Inhibiting POA^GAD2^→TMN neurons reduces REMs. **(A)** Left, schematic of optogenetic inhibition experiments. Center left, fluorescence image of POA in a GAD2-CRE mouse injected with AAVretro-DIO-SwiChR++-eYFP into the TMN. Scale bar, 1 mm. Center right, location of optic fiber tracts. Each colored bar represents the location of an optic fiber. Right, experimental paradigm for laser stimulation (2 s step pulses at 60 s intervals) in SwiChR++ and eYFP-expressing mice. **(B)** Example recording of a SwiChR++ (left) and eYFP mouse (right) with laser stimulation. Shown are EEG power spectra, EMG amplitude, and color-coded brain states. **(C)** Percentage of time spent in REMs, NREMs and wakefulness with and without laser in SwiChR++ and eYFP mice. Mixed ANOVA, virus p = 0.0629, laser p = 0.0003, interaction p = 0.0064; t-tests with Bonferroni correction, SwiChR-laser vs. SwiChR-w/o laser p = 3.00e-5, SwiChR-laser vs. eYFP-laser p = 0.0058. **(D)** Normalized EEG δ, θ, and σ power during REMs in SwiChR++ and eYFP mice with laser. Unpaired t-tests, SwiChR vs. eYFP p = 0.0432, 0.0099 for δ and σ. **(E)** Percentage of time spent in REMs, NREMs, and wakefulness during post laser sessions. **(C)** Bars, averages across mice; lines, individual mice; error bars, ± s.e.m. **(D, E)** Bars, averages across mice; dots, individual mice; error bars, ± s.e.m. SwiChR++: n = 12 mice; eYFP: n = 9 mice

Next, we compared the spectral composition of the EEG throughout recordings with or without laser stimulation in SwiChR++ and eYFP mice. During REMs, SwiChR++-mediated inhibition of POA^GAD2^→TMN neurons increased the δ and σ power in the EEG compared with SwiChR++-without laser and eYFP-laser groups (**Fig. 2D, S3C**). The δ, θ, and σ power in the EEG during NREMs and wake was indistinguishable between SwiChR++ and eYFP groups (**Fig. S3D**).

Next, we investigated the effect of sustained inhibition of POA^GAD2^→TMN neurons on the following 3 h recording without laser stimulation (**Fig. 2E**). Despite the reduction of REMs during the laser stimulation in SwiChR++ mice (**Fig. 2C**), there were no differences in the amount of REMs, NREMs, and wake between SwiChR++ and eYFP groups during the post laser recordings (**Fig. 2E, S3E, F**), suggesting that the loss of REMs during SwiChR++-mediated inhibition was not followed by a homeostatic increase in REMs.

In a complementary experiment, we also investigated how inhibition of TMN^HIS^ neurons regulates REMs using the same inhibitory optogenetics protocol. HDC-Cre mice were bilaterally injected with AAVs encoding Cre-inducible SwiChR++ (AAV_2_-EF1a-DIO-SwiChR++-eYFP) or eYFP (AAV_2_-Ef1α-DIO-eYFP) into the TMN followed by bilateral optic fibers implantation into the TMN (**Fig. S4A**). Consistent with the previous experiment, we compared SwiChR++ and eYFP recording with and without laser stimulation (**Fig. S4A, B**). SwiChR++-mediated inhibition of TMN^HIS^ neurons increased the amount of REMs compared with recordings without laser stimulation in the same mice and eYFP mice with laser stimulation (**Fig. S4C**). Given that TMN also contains other types of neurons, inhibition of histamine neurons by POA^GAD2^→TMN neurons may not be the sole source of the observed effect on REMs upon inhibition of POA^GAD2^→TMN neurons.

Taken together, we first demonstrated that SwiChR++-mediated inhibition of POA^GAD2^→TMN neurons reduced the amount of REMs, supporting a necessary role of these neurons in REMs regulation. Second, the lost amount of REMs was not compensated for in the subsequent sleep suggesting that their activity is involved in the homeostatic regulation of REMs.

### POA^GAD2^→TMN neurons exhibit an increased number of calcium transients during REMs restriction

To probe whether the activity of POA^GAD2^→TMN neurons changes during periods of high REMs pressure resulting from REMs restriction, we performed fiber photometry recordings combined with a closed-loop REMs restriction protocol (**Fig. 3A**). To detect the onset of REMs, we adapted a previously applied automatic REMs detection algorithm (Weber et al., 2015, 2018; Stucynski et al., 2022; Schott et al., 2023). Briefly, the animal’s brain state was classified based on real-time analysis of the EEG/EMG signals. As soon as a REMs episode was detected, a small vibrating motor attached to the animal’s head was turned on to terminate REMs by briefly awakening the animal (**Fig. 3A, S5A**) (Cardis et al., 2021; Osorio-Forero et al., 2023). Compared with the baseline recordings from the same mice during the same circadian time, we found that 6 h of REMs restriction (ZT 1.5-7.5) significantly reduced the amount of REMs and duration of REMs episodes, while increasing their frequency (**Fig. S5A, D-F;** Amount: Cohen’s d [d] = -1.621; Duration: d = -2.669; Frequency: d = 1.294). The number of motor activations gradually increased as mice tried to enter REMs more frequently, indicating the accumulation of REMs pressure (**Fig. S5B**). Before the onset of the motor vibration, we found a clear increase in the EEG θ band reflecting NREMs to REMs transitions (**Fig. S5C**). We also investigated the effects of REMs restriction on the spectral composition of the EEG. We found an increased δ power for REMs during restriction (**Fig. S5G**). During rebound (ZT 7.5-8.5), the amount of REMs was significantly increased compared with that during the same circadian time due to an increased frequency of REMs episodes (**Fig. S5H-K;** Amount: d = 0.868; Frequency: d = 1.147), compensating for the amount of REMs lost during restriction.

**FIGURE 3.**
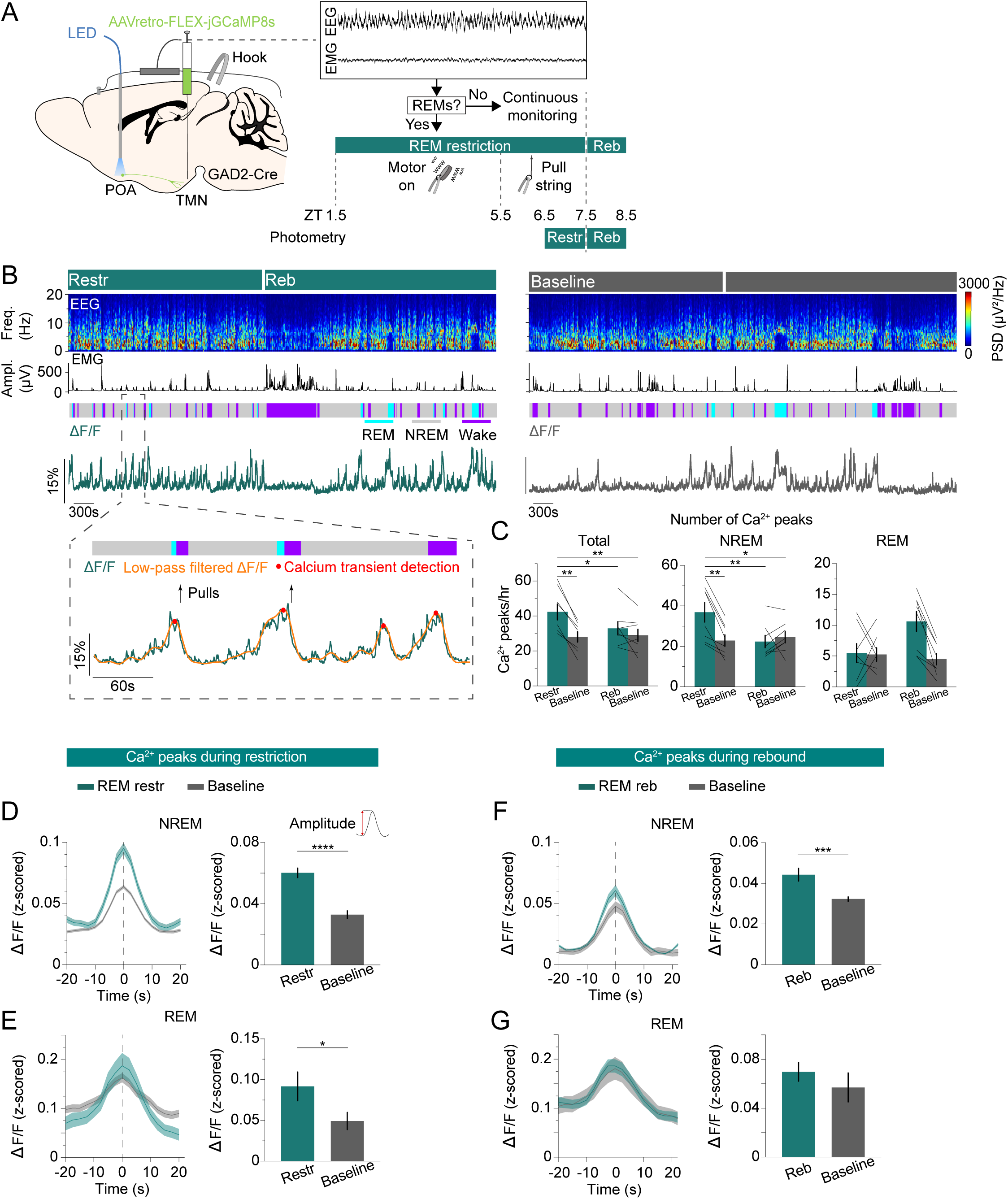
POA^GAD2^→TMN neurons exhibit an increased number of calcium transients during REMs restriction. **(A)** Schematic of REMs restriction/rebound and photometry recording experiments. The brain state was continuously monitored; once a REMs episode was detected, we used a vibrating motor (ZT 1.5-5.5) or pulled a string (ZT 5.5-7.5) attached to the mouse head to terminate REMs. Fiber photometry recordings were performed during REMs restriction (ZT 6.5-7.5) and rebound (ZT 7.5-8.5). **(B)** Top, example fiber photometry recording. Shown are EEG spectrogram, EMG amplitude, color-coded brain states, and ΔF/F signal. Bottom, brain states, ΔF/F signal (green), low-pass filtered ΔF/F signal (orange), detected peaks (red) and pulls (arrows) during a selected interval (dashed box) at an expanded timescale. **(C)** Number of calcium peaks during all states (left), NREMs (middle) and REMs (right). Bars, averages across mice; lines, individual mice; error bars, ± s.e.m. n = 8 mice. Total: two-way rm ANOVA, treatment (baseline vs. manipulation) p = 0.0031, time p = 0.0613, interaction p = 0.0189; t-tests with Bonferroni correction, REM restriction (restr.) vs. baseline (ZT 6.5-7.5) p = 0.0033, restr. vs. REM rebound (reb.) p = 0.0341, restr. vs. baseline (ZT 7.5-8.5) p = 0.0049. NREMs: two-way rm ANOVA, treatment p = 0.0233, time p = 0.0363, interaction p = 0.003; t-tests with Bonferroni correction, restr. vs. baseline (ZT 6.5-7.5) p = 0.0057, restr. vs. reb. p = 0.0046, restr. vs. baseline (ZT 7.5-8.5) p = 0.0115. **(D)** Left, average NREMs calcium peaks during REMs restriction and baseline recordings. Right, average amplitude of the NREMs calcium peaks. The amplitude was calculated by subtracting the ΔF/F value 10 s before the peak from its value at the peak. Unpaired t-tests, p = 6.30e-10. n = 295 and 196 peaks during restriction and baseline recordings. **(E)** Left, average REMs calcium peaks during REMs restriction and baseline recordings. Right, average amplitude of the REMs calcium peaks. Unpaired t-tests, p = 0.0437. n = 44 and 42 peaks during restriction and baseline recordings. **(F)** Left, average NREMs calcium peaks during REMs rebound and baseline recordings. Right, average amplitude of the NREMs calcium peaks. Unpaired t-tests, p = 0.0002. n = 180 and 196 peaks during rebound and baseline recordings. **(G)** Left, average REMs calcium peaks during REMs rebound and baseline recordings. Right, average amplitude of the REMs calcium peaks. n = 84 and 36 peaks during rebound and baseline recordings. **(D-G)** Bars, averages across trials; error bars, ± s.e.m; shadings, ± s.e.m.

During REMs restriction, we observed an increased number of calcium transients in the activity of POA^GAD2^→TMN neurons as the REMs pressure increased (**Fig. 3A, B**). To detect and quantify these transients, we applied an algorithm previously applied to detect calcium events in fiber photometry recordings (Antila et al., 2022; Smith et al., 2022). Using fiber photometry, we monitored the population activity of POA^GAD2^→TMN neurons during the last h of REMs restriction (ZT 6.5-7.5, referred as Restr.) and the first h of REMs rebound (ZT 7.5-8.5, referred as Reb.) and compared it with the activity during baseline recordings of the same mice on separate days at the same circadian time (**Fig. 3A, B**). To restrict REMs, we used a motor (ZT 1.5-5.5) as previously described or gently pulled a string attached to the animal’s head (ZT 5.5-7.5) during the photometry recordings (**Fig. 3A**). The manual REMs deprivation was utilized during the last 2 h of REMs restriction to avoid potential motion artifacts from the vibrating motor that could contaminate calcium signals. We verified that REMs was adequately restricted (**Fig. S6**). During restriction, the number of calcium transients of the POA^GAD2^→TMN neurons was significantly increased compared with that during baseline recordings (**Fig. 3C**). In particular, the number of calcium peaks was significantly elevated during NREMs, likely reflecting an increased pressure to transition to REMs (**Fig. 3C**). As the duration of REMs episodes was largely reduced as a result of the restriction (**Fig. S6A**), the number of peaks during REMs was not changed.

We further examined the amplitude of calcium transients during both NREMs and REMs. During REMs restriction, the amplitude of NREMs and REMs calcium transients was significantly higher compared with that during the circadian baseline (**Fig. 3D, E**) and remained elevated during NREMs in the rebound phase (**Fig. 3F, G**). Overall, these findings show that during periods of high REMs pressure, the POA^GAD2^→TMN neurons exhibit an increased number of calcium transients with higher amplitude compared with that during baseline levels suggesting that the activity of these neurons may reflect the heightened homeostatic need for REMs.

### Inhibition of POA^GAD2^→TMN neurons during REMs restriction reduces the REMs rebound

To investigate whether the activity of POA^GAD2^→TMN neurons encodes REMs pressure and consequently facilitates the subsequent rebound in REMs, we optogenetically inhibited these neurons during the last 3 h of REMs restriction. GAD2-Cre mice were bilaterally injected with retrograde AAVs encoding Cre-inducible SwiChR++ (AAVretro-DIO-SwiChR++-eYFP) or eYFP (AAVretro-DIO-eYFP) into the TMN followed by bilateral optic fiber implantation into the POA (**Fig. 4A**). Mice underwent REMs restriction (6 h, ZT 1.5-7.5), and laser stimulation (2 s step pulses at 60 s intervals) was applied during the last 3 h (ZT 4.5-7.5), when the REMs pressure was highest (**Fig. S4B**). During restriction, SwiChR-mediated inhibition of POA^GAD2^→TMN neurons decreased the percentage of REMs and reduced the frequency of REMs episodes with marginal significance (**Fig. 4B-D, S7A, B,** Amount: d = -1.254). We found that inhibition of POA^GAD2^→TMN neurons during REMs restriction resulted in a reduced amount of REMs during the rebound compared with that in eYFP mice (**Fig. 4B, F, G, S7D, E,** Amount: d = -1.317). Thus, inactivating POA^GAD2^→TMN neurons during heightened REMs pressure not only decreased the amount of REMs, but also prevented its homeostatic rebound during the following recovery sleep.

**FIGURE 4.**
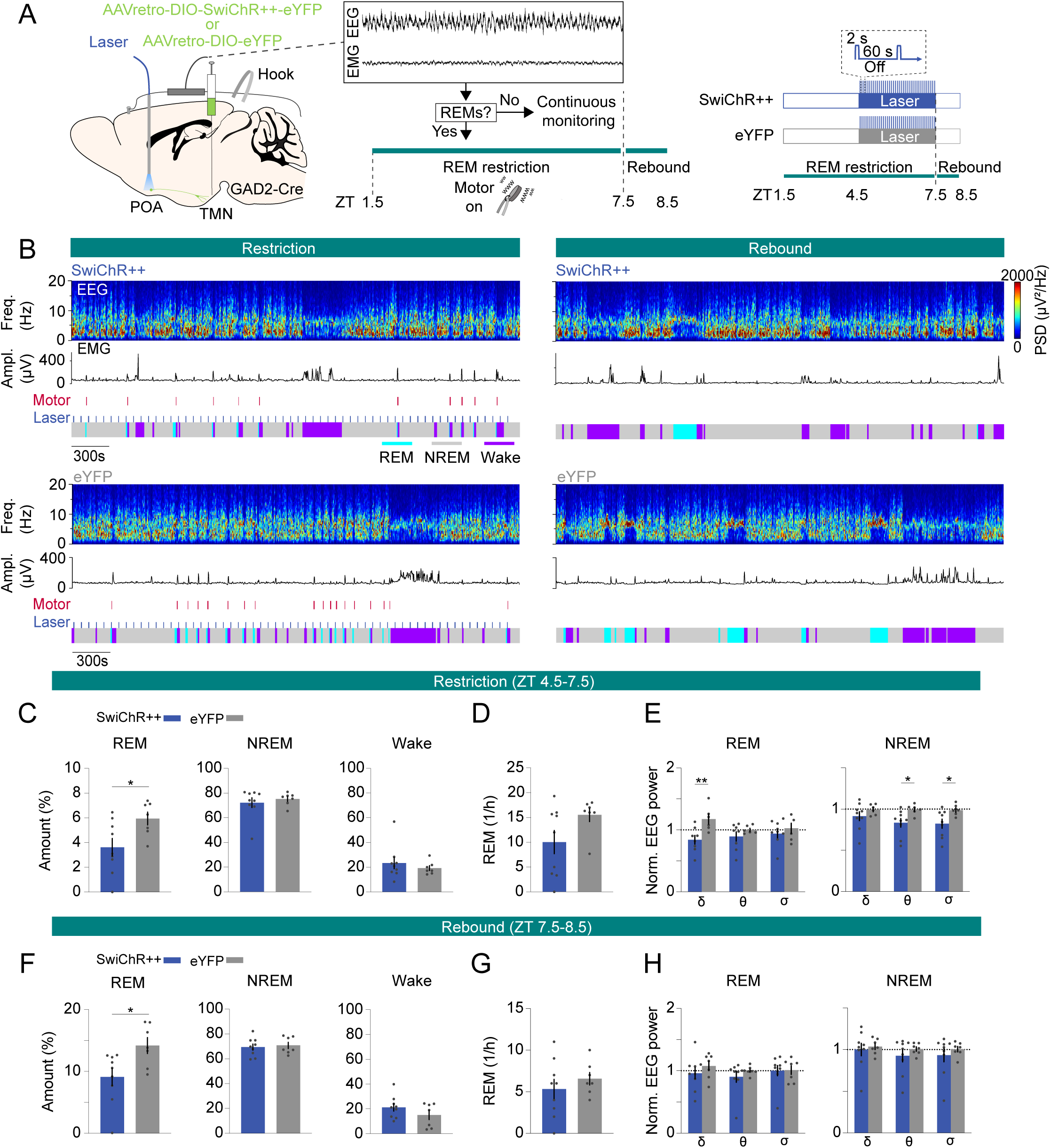
Inhibition of POA^GAD2^→TMN neurons during REMs restriction attenuates the REMs rebound. **(A)** Schematic of REMs restriction/rebound and optogenetic inhibition experiments. During closed-loop REMs restriction (ZT 1.5-7.5), a vibrating motor attached to the mouse head was used to terminate REMs. REMs was restricted for 6 h (ZT 1.5-7.5). During the last 3 h of restriction (ZT 4.5-7.5), laser stimulation (2 s step pulses at 60 s intervals) was applied in SwiChR++ and eYFP mice. **(B)** Example sessions from a SwiChR++ (top) and eYFP mouse (bottom) during REMs restriction with laser stimulation (left) and rebound (right). Shown are EEG spectrogram, EMG amplitude, motor vibration events, laser and color-coded brain states. **(C)** Percentage of time spent in REMs, NREMs and wakefulness during the last 3 h of REMs restriction with laser stimulation (ZT 4.5-7.5) in mice expressing SwiChR++ and eYFP. Unpaired t-tests, p = 0.026 for REMs amount. **(D)** Frequency of REMs episodes during the last 3 h of REMs restriction with laser stimulation (ZT 4.5-7.5). Unpaired t-tests, p = 0.0821. **(E)** Normalized EEG δ, θ and σ power during the last 3 h of REMs restriction with laser stimulation (ZT 4.5-7.5). Unpaired t-tests, p = 0.0091, 0.0332, and 0.038 for REMs δ, NREMs θ and σ power. **(F)** Percentage of time spent in REMs, NREMs and wakefulness during REMs rebound (ZT 7.5-8.5) in SwiChR++ and eYFP mice. Unpaired t-tests, p = 0.0205 for REMs amount. **(G)** Frequency of REMs episodes during REMs rebound (ZT 7.5-8.5). **(H)** Normalized EEG δ, θ and σ power during REMs rebound (ZT 7.5-8.5). Bars, averages across mice; dots, individual mice; error bars, ± s.e.m. SwiChR++: n = 9 mice; eYFP: n = 7 mice

Finally, we examined the spectral composition of the EEG during the REMs restriction. We found that inhibition of POA^GAD2^→TMN neurons during REMs restriction reduced the REMs δ power, NREMs θ and σ power compared with that in eYFP mice with laser stimulation (**Fig. 4E, S7C**). The reduced θ and σ power during NREMs could result from less attempts to enter REMs (**Fig. 4D**). During rebound, there were no significant differences in EEG δ, θ and σ power during REMs and NREMs (**Fig. 4H, S7F**).

Taken together, inhibiting POA^GAD2^→TMN neurons during REMs restriction significantly decreased the amount of REMs and blocked the subsequent rebound in REMs. Our data suggest that the heightened activity of POA^GAD2^→TMN neurons during sleep encodes the increased need for REMs and consequently plays an important role in the homeostatic response to REMs restriction.

## DISCUSSION

Our study demonstrates a role of POA^GAD2^→TMN neurons in the homeostatic regulation of REMs. Using fiber photometry, we showed that the POA^GAD2^→TMN neurons become activated throughout NREMs before transitioning to REMs, while being most active during REMs (**Fig. 1**). Sustained optogenetic inhibition of POA^GAD2^→TMN neurons reduced the overall amount of REMs, and the loss of REMs was not compensated during the subsequent recovery sleep (**Fig. 2**). During the period of high REMs pressure, POA^GAD2^→TMN neurons exhibited an increased number of calcium transients with elevated amplitude (**Fig. 3**). Optogenetic inhibition of POA^GAD2^→TMN neurons during REMs restriction attenuated the subsequent rebound of REMs (**Fig. 4**). Our results suggest that the activity of POA^GAD2^→TMN neurons reflects an increased need for REMs in the form of enhanced calcium transients and is required for the rebound following the loss of REMs.

The TMN contains histamine-producing neurons and antagonizing histamine signaling causes sleepiness (Watanabe et al., 1983; Panula et al., 1984; Lin et al., 1988; Bayliss et al., 1990; Haas et al., 2008; Uygun et al., 2016). Consistent with our photometry recordings (**Fig. S2**), electrophysiological recordings demonstrate that TMN^HIS^ neurons are most active during wakefulness and less active during NREMs and REMs (Steininger et al., 1999; Vanni-Mercier et al., 2003; John et al., 2004; Takahashi et al., 2006). Throughout NREMs, the activity of TMN^HIS^ neurons gradually decreased while that of POA^GAD2^→TMN neurons showed an opposite pattern (**Fig 1E, S2E**), which is likely in part the result of direct synaptic inputs from the POA^GAD2^→TMN neurons to TMN^HIS^ neurons (Chung et al., 2017; Saito et al., 2018). TMN^HIS^ neurons in turn inhibit putative sleep-active POA neurons (Williams et al., 2014). Mutual inhibition between TMN^HIS^ and POA^GAD2^→TMN neurons may explain their antagonistic activity pattern revealed in our fiber photometry recordings.

While many studies have focused on the role of the POA in regulating NREMs (Sherin et al., 1996; Szymusiak et al., 1998; Lu et al., 2000; Zhang et al., 2015; Kroeger et al., 2018; Ma et al., 2019), previous *in vivo* electrophysiological studies found that the majority of sleep-active neurons in the POA are most active during REMs (Osaka and Matsumura, 1995; Takahashi et al., 2009; Antila et al., 2022). Similarly, recent fiber photometry recordings demonstrated that GABAergic neurons in the POA and their subtypes expressing cholecystokinin, corticotropin-releasing hormone, tachykinin 1 or galanin are most active during REMs (Miracca et al., 2022; Smith et al., 2022). Deleting the NMDA receptor GluN1 subunit in the POA reduced REMs (Miracca et al., 2022), and in line with this, sustained optogenetic inhibition of POA^GAD2^→TMN neurons specifically decreased REMs. Consistent with these studies, our findings also support an important role of the POA in REMs regulation and, in addition, provide evidence that these neurons are also part of the homeostat regulating REMs.

A previous study showed that the number of c-Fos positive POA neurons is positively correlated with the amount of REMs in rats (Lu et al., 2002). Moreover, REMs restriction led to an increase in the number of POA neurons expressing c-Fos, which was correlated with the number of attempts to enter REMs (Gvilia et al., 2006). Together with our findings that the number of calcium transients of POA^GAD2^→TMN neurons increased during REMs restriction and that inhibition of these neurons blocked the following REMs rebound, these results support a crucial role of the POA in the homeostatic regulation of REMs. For the future, it would be interesting to test whether POA neurons projecting to other postsynaptic areas are also involved in the homeostatic regulation of REMs. A previous study showed that POA neurons projecting to REMs-regulatory pontine regions including the laterodorsal tegmental nucleus, locus coeruleus (LC) and dorsal raphe nucleus express increased levels of c-Fos after periods of dark exposure that increased REMs (Lu et al., 2002). However, the number of c-Fos positive POA neurons projecting to the LC was not increased upon REMs restriction, suggesting that this subpopulation may not be involved in the homeostatic regulation of REMs (Verret et al., 2006). Besides the TMN, the POA also projects to other REMs-regulatory regions such as the ventrolateral periaqueductal gray (vlPAG) and lateral hypothalamus (Steininger et al., 2001; Saito et al., 2018). Particularly, the projections to the vlPAG are of interest for future research, as GABAergic neurons in this area have been previously implicated in the homeostatic regulation of REMs (Hayashi et al., 2015; Weber et al., 2018). It remains to be tested whether POA^GAD2^→TMN neurons also project to these brain regions to potentially regulate REMs homeostasis.

The cellular mechanisms underlying the elevated activity of POA neurons during high REMs pressure are unknown. Sleep-promoting neurons in the dorsal fan-shaped body of *Drosophila* display increased intrinsic neuronal excitability in response to sleep need (Donlea et al., 2014). REMs deprivation was shown to change the intrinsic excitability of hippocampal neurons and impact synaptic plasticity (McDermott et al., 2003; Mallick and Singh, 2011; Zhou et al., 2020). The elevated activity of POA^GAD2^→TMN neurons during heightened REMs pressure may similarly be the result of an increased excitability. Given that the POA is also involved in the homeostatic regulation of NREMs (Sherin et al., 1996; Szymusiak et al., 1998; Gong et al., 2004; Zhang et al., 2015; Ma et al., 2019; Lu et al., 2000), it would be interesting to study how different POA subpopulations integrate the homeostatic need for NREMs and REMs.

Together, we have demonstrated a role of POA^GAD2^→TMN neurons in the homeostatic regulation of REMs. REMs disturbances are observed in a variety of psychiatric disorders such as depression and PTSD and often precede their clinical onset (Ross et al., 1989; Gottesmann and Gottesman, 2007; Germain, 2013; Palagini et al., 2013). Elucidating the circuit mechanisms underlying the homeostatic regulation of REMs may provide novel therapeutic targets to specifically regulate and normalize REMs in these psychiatric disorders to alleviate associated symptoms and potentially slow down their progression.

## MATERIALS and METHODS

### Mice

All experimental procedures were approved by the Institutional Animal Care and Use Committee (IACUC reference # 806197) at the University of Pennsylvania and conducted in compliance with the National Institutes of Health Office of Laboratory Animal Welfare Policy. Experiments were performed in male and female GAD2-IRES-Cre mice (#010802, Jackson Laboratory, generously donated by Taniguchi et al., 2011) or HDC-IRES-Cre mice (#021198, Jackson Laboratory, generously donated by Zecharia et al., 2012) aged 10-18 weeks, weighing 18-25 g at the time of surgery. Animals were group-housed with littermates on a 12h light/12h dark cycle (lights on 7 am and off 7 pm) with *ad libitum* access to food and water.

### Viruses

Cre-dependent adeno-associated viral vectors were used to selectively express GCaMP, SwiChR++ or eYFP in POA^GAD2^ →TMN neurons, POA^GAD2^ →TMN projections or TMN^HIS^ neurons. pGP-AAV-Syn-FLEX-jGCaMP8s-WPRE was developed from the GENIE Project (Zhang et al., 2023) (162377-AAVrg, Addgene). pAAV-Syn-FLEX-GCaMP6s-WPRE-SV40 was developed from Douglas Kim & GENIE Project (Chen et al., 213) (Penn Vector Core or 100845-AAV1, Addgene). rAAV_2_-Retro–Ef1α-DIO-SwiChR++-eYFP (R47730, UNC Vector Core). rAAV_2_-Retro–Ef1α-DIO-eYFP (R49556, UNC Vector Core).

### Surgical procedures

All procedures followed the IACUC guidelines for rodent survival surgery. Mice were anesthetized with isoflurane (1-2%) during the surgery, and placed on a stereotaxic frame (Kopf) while being on a heating pad to maintain body temperature. The skin was incised and small holes were drilled for virus injections and implantations of optic fibers and EEG/EMG electrodes.

For fiber photometry experiments to image POA^GAD2^→TMN neurons (**Fig. 1, 3**), pGP-AAV-Syn-FLEX-jGCaMP8s-WPRE was injected (Nanoject II, Drummond Scientific) into the TMN (300 nl; AP -2.4 mm; ML -1 mm; DV -5.4 to -5.2 mm, relative to bregma) and an optic fiber (400 μm diameter) was implanted into the POA (AP 0.2 mm; ML -0.6 mm; DV -5.2 mm). For imaging the POA^GAD2^ →TMN axonal fibers (**Fig. S1**), pAAV-Syn-FLEX-GCaMP6s-WPRE-SV40 was injected into the POA (300 nl) and an optic fiber was implanted into the TMN. To image TMN^HIS^ neurons (**Fig. S2**), pAAV-Syn-FLEX-GCaMP6s-WPRE-SV40 was injected into the TMN (300 nl) and an optic fiber was implanted into the TMN.

For optogenetic inhibition experiments (**Fig. 2, 4**), rAAV_2_-Retro–Ef1α-DIO-SwiChR++-eYFP (for inhibition group) or rAAV_2_-Retro–Ef1α-DIO-eYFP (for control group) was bilaterally injected into the location on the reference picture the number of mice with overlapping virus expression, which was encoded using different green color intensities.

### Polysomnographic recordings

All sleep recordings were performed in a cage to which the animal had been habituated for several days. All recordings were performed during the light phase between 8 am and 5 pm (ZT1-10) in sound-attenuating chambers. For sleep recordings, EEG and EMG signals were recorded using an RHD2132 amplifier (Intan Technologies, sampling rate 1 kHZ) connected to an RHD USB interface board (Intan Technologies). For fiber photometry experiments, a calcium signal was recorded using an RZ5P amplifier (Tucker-Davis Technologies, sampling rate 1.5 kHz). EEG and EMG signals were referenced to a ground screw located on top of the cerebellum. At the start of each sleep recording, EEG and EMG electrodes were connected to flexible recording cables via small connectors. To determine the brain state of the animal, we first computed the EEG and EMG spectrogram for sliding, half-overlapping 5 s windows, resulting in 2.5 s time resolution. To estimate within each 5 s window the power spectral density (PSD), we performed Welch’s method with Hanning window using sliding, half-overlapping 2 s intervals. Next, we computed the time-dependent δ (0.5 to 4 Hz), θ (5 to 12 Hz), σ (12 to 20 Hz), and high γ (100 to 150 Hz) power by integrating the EEG power in the corresponding ranges within the EEG spectrogram. In addition, we calculated the ratio of the θ and δ power (θ/δ) and the EMG power in the range of 50 to 500 Hz. For each power band, we used its temporal mean to separate it into a low and high part (except for the EMG and θ/δ ratio, where we used the mean plus one standard deviation as threshold). REMs was defined by a high θ/δ ratio, low EMG, and low δ power. NREMs was defined by high δ power, a low θ/δ ratio, and low EMG power. In addition, states with low EMG power, low δ power, but high σ power were scored as NREMs. Wake was defined by low δ power, high EMG power and high γ power (if not otherwise classified as REMs). Our automatic algorithm that has been previously used in (Weber et al., 2015, 2018; Chung et al., 2017; Antila et al., 2022; Smith et al., 2022; Stucynski et al., 2022; Schott et al., 2023) has 90.256 % accuracy compared with the manual scoring by expert annotators. We manually verified the automatic classification using a graphical user interface visualizing the raw EEG and EMG signals, EEG spectrograms, EMG amplitudes, and the hypnogram to correct for errors, by visiting each single 2.5 s epoch in the hypnograms. The software for automatic brain state classification and manual scoring was programmed in Python (https://github.com/tortugar/Lab/tree/master/PySleep).

### Fiber photometry

Prior to the recording, the optic fiber and EEG/EMG electrodes were connected to flexible patch cables. For calcium imaging, a first LED (Doric lenses) generated the excitation wavelength of 465 nm and a second LED emitted 405 nm light, which served as control for bleaching and motion artifacts. 465 and 405 nm signals were modulated at two different frequencies, 210 and 330 Hz respectively. Both lights traveled through dichroic mirrors (Doric lenses) before entering a patch cable attached to the optic fiber. Fluorescence signals emitted by GCaMP8s or GCaMP6s were collected by the optic fiber and traveled via the patch cable through a dichroic mirror and GFP emission filter (Doric lenses) before entering a photoreceiver (Newport Co.). Photoreceiver signals were relayed to an RZ5P amplifier and demodulated into two signals using the Synapse software (Tucker-Davis Technologies), corresponding to the 465 and 405 nm excitation wavelengths. To analyze the calcium activity, we used custom-written Python scripts. First, both signals were low-pass filtered at 2 Hz using a 4th order digital Butterworth filter. Next, we fitted the 405 nm to the 465 nm signal using linear regression. Finally, the linear fit was subtracted from the 465 nm signal to correct for photobleaching and/or motion artifacts, and the difference was divided by the linear fit yielding the ΔF/F signal. Both the fluorescence signals and EEG/EMG signals were simultaneously recorded using the RZ5P amplifier. Fiber photometry recordings were excluded if the signals suddenly shifted, likely due to a loose connection between the optic fiber and patch cable.

### Optogenetic manipulation

Sleep recordings were performed during the light phase (ZT2-8). Mice were tethered to bilateral patch cables connected with the lasers and a flexible recording cable to record EEG/EMG signals. The recording started after 30 min of habituation. For optogenetic inhibition experiments, 2 s step pulses (1-3 mW, 60 s intervals) were generated by a blue laser (473 nm, Laserglow) and sent through the optic fiber (200 um diameter, Thorlabs) connected to the ferrule on the animal’s head for 3 h (ZT2-5). This laser stimulation protocol was rationally designed based on previous reports of sustained inhibition and prior results that recapitulate similar findings as inhibitory chemogenetic techniques (Iyer et al., 2016; Kim et al., 2016; Wiegert et al., 2017; Stucynski et al., 2022). TTL pulses to trigger the laser were controlled using a Raspberry Pi, which was controlled by a custom-programmed user interface programmed in Python. Following sustained inhibition, an additional 3 h (ZT5-8) recording was performed without laser stimulation (post-laser session). Baseline recordings were performed (without laser stimulation, ZT2-5), and counterbalanced across mice and days to avoid potential order effects. For each mouse, we collected 2-3 baseline and laser recordings each.

### REMs restriction

We employed an automatic REMs detection algorithm (Weber et al., 2015, 2018; Stucynski et al., 2022; Schott et al., 2023) and used a small vibrating motor to terminate/restrict REMs (Cardis et al., 2021; Osorio-Forero et al., 2023) (https://github.com/luthilab/IntanLuthiLab). A small vibrating motor (DC 3V Mini Vibration Motor, diameter: 10mm, thickness: 3mm, BOJACK) with a soldered lobster claw clasp was attached to a small wire hook secured in the dental cement of each mouse head. The motors had cables that were connected to a Raspberry Pi which controlled the motor onset and offset.

The automatic REMs detection algorithm determined whether the mouse was in REMs or not based on real-time spectral analysis of the EEG/EMG signals. The onset of REMs was defined as the time point where the EEG θ/δ ratio exceeded a threshold (mean + 1 std of θ/δ), which was calculated from the same mouse using previous recordings. As soon as a REMs episode was detected, the small vibrating motor turned on to terminate REMs and consequently woke up the mouse. The motor vibrated until the REMs episode was terminated, i.e. when the θ/δ ratio dropped below its mean value or if the EMG amplitude surpassed a threshold (mean + 0.5 std of amplitude). All REMs restriction experiments started at ZT1.5 and lasted until ZT7.5. Following the restriction, motors were turned off and mice were permitted to enter recovery sleep.

To monitor the calcium activity during REMs restriction using fiber photometry, we performed manual REMs restriction to avoid potential motion artifacts caused by the vibrating motor that could interfere with the integrity of the fiber photometry signals. Briefly mice underwent a 4 h automatic REMs restriction protocol as described above. During the last 2 h of restriction, REMs was detected manually based on the EEG and EMG signal by an experimenter who gently pulled on a string attached to the hook secured in the dental cement of the mouse head. To avoid potential photobleaching, the recording was performed during the last 1 h of REMs restriction and 1 h of REMs rebound. Each mouse also underwent a baseline recording on a separate day. The order of REMs restriction and baseline recordings were varied to minimize the impact of the experimental sequence on the results.

We also performed automatic REMs restriction combined with optogenetic inhibition. Each mouse underwent REMs restriction for 6 h, and we continuously delivered 2 s step pulses (1-3 mw, 473 nm, Laserglow) at 60 s intervals during the last 3 h of restriction (ZT4.5-7.5). Following REMs restriction, the recovery sleep was recorded.

### Analysis of ΔF/F activity at brain state transitions and during NREMs

To calculate the neural activity changes relative to brain state transitions (**Fig. 1D, Fig. S1D, Fig. S2D**), we aligned the ΔF/F signals for transitions across all mice relative to the time point of the transition (t = 0 s). For each NREMs→ REMs or NREMs→Wake transition, we ensured that the preceding NREMs episodes lasted for at least 60 s, only interrupted by short awakenings (≤ 20 s). To determine the time point at which the activity significantly started to increase or decrease, we used the first 10 s of NREMs as baseline. For REMs→Wake and Wake→NREMs transitions, the preceding REMs or Wake episode was at least 30 s long. Using one-way rm ANOVA, we tested whether the activity (downsampled to 10s bins) within each 10 s bin was significantly modulated throughout the transition (from -60 to 30 s). Finally, using pairwise t-tests with Holm-Bonferroni correction, we determined the time bins for which the activity significantly differed from the baseline bin (activity for bin -60 to -50 s). The time point for a given 10 s bin was set to its midpoint. To account for multiple comparisons, we divided the significance level (α = 0.05) by the number of comparisons (Bonferroni correction). To analyze the activity throughout NREMs, we normalized the duration of all NREMs episodes and the corresponding ΔF/F signals to the same length (**Fig. 1E, Fig. S2DE**).

### Spectrotemporal correlation analysis

To identify features of the EEG spectrogram associated with POA^GAD2^→TMN calcium activity, we adapted a receptive field model (Weber et al., 2010; Schott et al., 2023) to predict POA^GAD2^→TMN neural activity from the spectrogram. Intuitively, we estimated a spectrotemporal filter using linear regression that predicts for each time point the POA^GAD2^ →TMN calcium response. In more detail, we first computed the EEG spectrogram *E(f_i_, t_j_)* using 2 s windows with 80 % overlap, resulting in a time resolution *dt* of 400 ms. Each spectrogram frequency was normalized by its mean power across the recording, and the parameter *E(f_i_, t_j_)* specifies the relative amplitude of frequency *f_i_* for time point *t_j_*. We then downsampled the ΔF/F response using the same time resolution as for the spectrogram, and extracted all time bins with REMs, NREMs or wake for analysis.

To predict the calcium response, the EEG spectrogram was linearly filtered with the kernel *H(f_i_, t_l_)*. In analogy to receptive fields estimated using similar approaches for sensory neurons, we used the term “spectral field” for *H(f_i_, t_l_)*. The spectral field can be described as the set of coefficients that optimally relate the neural activity at time *t_j_* to the EEG spectrogram at time *t_j + l_*, where *l* represents the lag between the time points of the spectrogram and the neural response. The estimated frequency components of *H(f_i_, t_l_)* ranged from *f_1_* = 0.5 Hz to *f_nf_* = 20 Hz, while the time axis ranged from -*n_T_* * *dt* = -50 s to *n_T_*\* *dt* = 50 s. Thus, *H(f_i_, t_l_)* comprises *n_f_* frequencies and 2 * *n_T_*+ 1 time bins over a window from -50 to +50 seconds relative to the neural response. The convolution of the EEG spectrogram with the spectral field can be expressed as

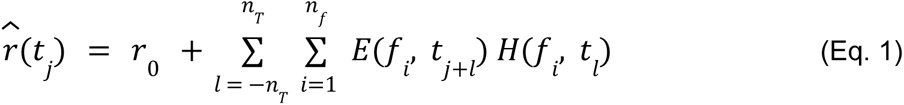

The scalar parameter *r_0_* denotes a constant offset, and *r̂*(*t_j_*) denotes the predicted neural response. The optimal spectral field *H(f_i_, t_l_)* minimizes the mean-squared error between the predicted and measured neural response at time *t_j_*. To account for the large number of estimated parameters, we included a regularization term in the error function, which penalizes large kernel components and therefore guards against overfitting of the model. The spectral field for each recording was estimated using 5-fold cross-validation to determine the regularization parameter (λ) that optimized the average model performance on the test sets. Kernels were averaged across all recordings for individual animals; the spectrotemporal correlation in **Fig. 1F** represents the mean spectral field across all mice.

### Power spectral density and power estimation

The power spectral density (PSD) of the EEG was computed using Welch’s method with the Hanning window for sliding, half-overlapping 2 s intervals. To calculate the power within a given frequency band, we approximated the corresponding area under the spectral density curve using a midpoint Riemann sum. To compute the EMG amplitude, we calculated the PSD of the EMG and integrated frequencies between 5 - 100 Hz. To test whether laser stimulation changed the spectral density during a specific brain state, we determined for each mouse the δ, θ or σ power for that state with and without laster. To compare the PSD between experimental and control mice, we calculated the δ, θ, and σ power with and without laser stimulation for each mouse and normalized the power values with laser by the power obtained for epochs without laser.

### Detection of calcium transients

To detect calcium transients, we first filtered the ΔF/F signal with a zero-lag, 4th order digital Butterworth filter with cutoff frequency of 1/20 Hz. Next, prominent peaks in the signal were detected using the function scipy.find_peaks provided by the open source Python library scipy (https://scipy.org). As parameter for the peak prominence, we used 0.15 * distance between the 1st and 99th percentile of the distribution of the ΔF/F signal. A transient was defined as occurring during NREMs and REMs, if the peak overlapped with NREMs or REMs respectively. The calcium transient amplitude was calculated by using the values 10 s preceding the peak and subtracting that from the peak values at 0 s.

### Statistical tests

Statistical analyses were performed using the python modules (scipy.stats, https://scipy.org; pingouin, https://pingouin-stats.org) and Prism v9.5.0.0 (GraphPad Software Inc). We did not predetermine sample sizes, but cohorts were similarly sized as in other relevant sleep studies (Jego et al., 2013; Ma et al., 2019). All data collection was randomized and counterbalanced. All data are reported as mean ± s.e.m. A (corrected) p-value < 0.05 was considered statistically significant for all comparisons. Data were compared using unpaired t-tests, paired t-tests, one-way ANOVAs, or two-way ANOVAs followed by multiple comparisons as appropriate. The statistical results for the figures are presented in the Table S1, and Figure legends.

## Data sharing plans

The code used for data analysis is publicly available under: https://github.com/tortugar/Lab. All the data are available from the corresponding author upon reasonable request.

## Acknowledgements

We thank Mandy Schott for generating python scripts for optogenetic manipulation during REMs restriction, Jenny Smith for help with generating the virus expression heatmaps and members of the Chung and Weber labs for helpful discussion. This work was funded by the National Institute of Neurological Disorders and Stroke (R01-NS-110865), the Whitehall Foundation, the Alfred P. Sloan Foundation, a NARSAD Young Investigator Grant from the Brain & Behavior Research Foundation, a Simons Foundation Pilot Award, an Eagle Autism Challenge Pilot Grant, the Thomas B. and Jeannette E. Laws McCabe Fund Award, the Hartwell Individual Biomedical Research Award (to S.C.), and the NIH individual F31 fellowship (NS118963-01A1, to J.M).

## Author contributions

Conceptualization: J.M., F.W., and S.C.; Methodology: J.M., X.J., J.H., R.C., A.O-F. A.Lüthi, F.W., and S.C.; Software: J.M., X.J., and F.W.; Validation: J.M., A.Lin, and N.S.; Formal Analysis: J.M.; Investigation: J.M. and A.Lin; Resources: J.M., F.W., and S.C.; Data Curation: J.M. and A.Lin; Writing –Original Draft: J.M.; Writing –Review and Editing: J.M., R.C., A.O-F. A.Lüthi, F.W., and S.C.; Visualization: J.M.; Supervision: S.C.; Funding Acquisition: J.M. and S.C.

## REFERENCES

Antila, H., Kwak, I., Choi, A., Pisciotti, A., Covarrubias, I., Baik, J., et al. (2022). A noradrenergic-hypothalamic neural substrate for stress-induced sleep disturbances. Proc. Natl. Acad. Sci. U. S. A. 119, e2123528119. doi: 10.1073/pnas.2123528119.

Bayliss, D. A., Wang, Y. M., Zahnow, C. A., Joseph, D. R., and Millhorn, D. E. (1990). Localization of histidine decarboxylase mRNA in rat brain. Mol. Cell. Neurosci. 1, 3–9. doi: 10.1016/1044-7431(90)90036-4.

Beersma, D. G., Dijk, D. J., Blok, C. G., and Everhardus, I. (1990). REM sleep deprivation during 5 hours leads to an immediate REM sleep rebound and to suppression of non-REM sleep intensity. Electroencephalogr. Clin. Neurophysiol. 76, 114–122. doi: 10.1016/0013-4694(90)90209-3.

Benington, J. H., Woudenberg, M. C., and Heller, H. C. (1994). REM-sleep propensity accumulates during 2-h REM-sleep deprivation in the rest period in rats. Neurosci. Lett. 180, 76–80. doi: 10.1016/0304-3940(94)90917-2.

Berndt, A., Lee, S. Y., Wietek, J., Ramakrishnan, C., Steinberg, E. E., Rashid, A. J., et al. (2016). Structural foundations of optogenetics: Determinants of channelrhodopsin ion selectivity. Proc. Natl. Acad. Sci. U. S. A. 113, 822–829. doi: 10.1073/pnas.1523341113.

Cardis, R., Lecci, S., Fernandez, L. M., Osorio-Forero, A., Chu Sin Chung, P., Fulda, S., et al. (2021). Cortico-autonomic local arousals and heightened somatosensory arousability during NREMS of mice in neuropathic pain. eLife 10, e65835. doi: 10.7554/eLife.65835.

Chen, T.-W., Wardill, T. J., Sun, Y., Pulver, S. R., Renninger, S. L., Baohan, A., et al. (2013). Ultrasensitive fluorescent proteins for imaging neuronal activity. Nature 499, 295–300. doi: 10.1038/nature12354.

Chung, S., Weber, F., Zhong, P., Tan, C. L., Nguyen, T. N., Beier, K. T., et al. (2017). Identification of preoptic sleep neurons using retrograde labelling and gene profiling. Nature 545, 477–481. doi: 10.1038/nature22350.

Clément, O., Sapin, E., Bérod, A., Fort, P., and Luppi, P.-H. (2011). Evidence that neurons of the sublaterodorsal tegmental nucleus triggering paradoxical (REM) sleep are glutamatergic. Sleep 34, 419–423. doi: 10.1093/sleep/34.4.419.

Donlea, J. M., Pimentel, D., and Miesenböck, G. (2014). Neuronal Machinery of Sleep Homeostasis in Drosophila. Neuron 81, 860–872. doi: 10.1016/j.neuron.2013.12.013.

Endo, T., Schwierin, B., Borbély, A. A., and Tobler, I. (1997). Selective and total sleep deprivation: effect on the sleep EEG in the rat. Psychiatry Res. 66, 97–110. doi: 10.1016/s0165-1781(96)03029-6.

Endo, T., Roth, C., Landolt, H. P., Werth, E., Aeschbach, D., Achermann, P., et al. (1998). Selective REM sleep deprivation in humans: effects on sleep and sleep EEG. Am. J. Physiol. 274, R1186–1194. doi: 10.1152/ajpregu.1998.274.4.R1186.

Franken, P. (2002). Long-term vs. short-term processes regulating REM sleep. J. Sleep Res. 11, 17–28. doi: 10.1046/j.1365-2869.2002.00275.x.

Germain, A. (2013). Sleep disturbances as the hallmark of PTSD: where are we now? Am. J. Psychiatry 170, 372–382. doi: 10.1176/appi.ajp.2012.12040432.

Gong, H., McGinty, D., Guzman-Marin, R., Chew, K.-T., Stewart, D., and Szymusiak, R. (2004). Activation of c-fos in GABAergic neurones in the preoptic area during sleep and in response to sleep deprivation. J. Physiol. 556, 935–946. doi: 10.1113/jphysiol.2003.056622.

Gottesmann, C. (1996). The transition from slow-wave sleep to paradoxical sleep: evolving facts and concepts of the neurophysiological processes underlying the intermediate stage of sleep. Neurosci. Biobehav. Rev. 20, 367–387. doi: 10.1016/0149-7634(95)00055-0.

Gottesmann, C., and Gottesman, I. (2007). The neurobiological characteristics of rapid eye movement (REM) sleep are candidate endophenotypes of depression, schizophrenia, mental retardation and dementia. Prog. Neurobiol. 81, 237–250. doi: 10.1016/j.pneurobio.2007.01.004.

Gvilia, I., Turner, A., McGinty, D., and Szymusiak, R. (2006). Preoptic Area Neurons and the Homeostatic Regulation of Rapid Eye Movement Sleep. J. Neurosci. 26, 3037–3044. doi: 10.1523/JNEUROSCI.4827-05.2006.

Haas, H. L., Sergeeva, O. A., and Selbach, O. (2008). Histamine in the Nervous System. Physiol. Rev. 88, 1183–1241. doi: 10.1152/physrev.00043.2007.

Hayashi, Y., Kashiwagi, M., Yasuda, K., Ando, R., Kanuka, M., Sakai, K., et al. (2015). Cells of a common developmental origin regulate REM/non-REM sleep and wakefulness in mice. Science 350, 957–961. doi: 10.1126/science.aad1023.

Iyer, S. M., Vesuna, S., Ramakrishnan, C., Huynh, K., Young, S., Berndt, A., et al. (2016). Optogenetic and chemogenetic strategies for sustained inhibition of pain. Sci. Rep. 6, 30570. doi: 10.1038/srep30570.

Jego, S., Glasgow, S. D., Herrera, C. G., Ekstrand, M., Reed, S. J., Boyce, R., et al. (2013). Optogenetic identification of a rapid eye movement sleep modulatory circuit in the hypothalamus. Nat. Neurosci. 16, 1637–1643. doi: 10.1038/nn.3522.

John, J., Wu, M.-F., Boehmer, L. N., and Siegel, J. M. (2004). Cataplexy-Active Neurons in the Hypothalamus: Implications for the Role of Histamine in Sleep and Waking Behavior. Neuron 42, 619–634. doi: 10.1016/S0896-6273(04)00247-8.

Kim, H., Ährlund-Richter, S., Wang, X., Deisseroth, K., and Carlén, M. (2016). Prefrontal Parvalbumin Neurons in Control of Attention. Cell 164, 208–218. doi: 10.1016/j.cell.2015.11.038.

Kroeger, D., Absi, G., Gagliardi, C., Bandaru, S. S., Madara, J. C., Ferrari, L. L., et al. (2018). Galanin neurons in the ventrolateral preoptic area promote sleep and heat loss in mice. Nat. Commun. 9, 4129. doi: 10.1038/s41467-018-06590-7.

Lin, J.-S., Sakai, K., and Jouvet, M. (1988). Evidence for histaminergic arousal mechanisms in the hypothalamus of cat. Neuropharmacology 27, 111–122. doi: 10.1016/0028-3908(88)90159-1.

Lu, J., Greco, M. A., Shiromani, P., and Saper, C. B. (2000). Effect of lesions of the ventrolateral preoptic nucleus on NREM and REM sleep. J. Neurosci. Off. J. Soc. Neurosci. 20, 3830–3842. doi: 10.1523/JNEUROSCI.20-10-03830.2000.

Lu, J., Bjorkum, A. A., Xu, M., Gaus, S. E., Shiromani, P. J., and Saper, C. B. (2002). Selective activation of the extended ventrolateral preoptic nucleus during rapid eye movement sleep. J. Neurosci. Off. J. Soc. Neurosci. 22, 4568–4576. doi: 10.1523/JNEUROSCI.22-11-04568.2002.

Ma, Y., Miracca, G., Yu, X., Harding, E. C., Miao, A., Yustos, R., et al. (2019). Galanin Neurons Unite Sleep Homeostasis and α2-Adrenergic Sedation. Curr. Biol. CB 29, 3315–3322.e3. doi: 10.1016/j.cub.2019.07.087.

Mallick, B. N., and Singh, A. (2011). REM sleep loss increases brain excitability: role of noradrenaline and its mechanism of action. Sleep Med. Rev. 15, 165–178. doi: 10.1016/j.smrv.2010.11.001.

McDermott, C. M., LaHoste, G. J., Chen, C., Musto, A., Bazan, N. G., and Magee, J. C. (2003). Sleep deprivation causes behavioral, synaptic, and membrane excitability alterations in hippocampal neurons. J. Neurosci. Off. J. Soc. Neurosci. 23, 9687–9695. doi: 10.1523/JNEUROSCI.23-29-09687.2003.

McGinty, D. J., and Sterman, M. B. (1968). Sleep suppression after basal forebrain lesions in the cat. Science 160, 1253–1255. doi: 10.1126/science.160.3833.1253.

Miracca, G., Anuncibay-Soto, B., Tossell, K., Yustos, R., Vyssotski, A. L., Franks, N. P., et al. (2022). NMDA Receptors in the Lateral Preoptic Hypothalamus Are Essential for Sustaining NREM and REM Sleep. J. Neurosci. Off. J. Soc. Neurosci. 42, 5389–5409. doi: 10.1523/JNEUROSCI.0350-21.2022.

Nauta, W. J. H. (1946). Hypothalamic regulation of sleep in rats; an experimental study. J. Neurophysiol. 9, 285–316. doi: 10.1152/jn.1946.9.4.285.

Osaka, T., and Matsumura, H. (1995). Noradrenaline inhibits preoptic sleep-active neurons through alpha 2-receptors in the rat. Neurosci. Res. 21, 323–330. doi: 10.1016/0168-0102(94)00871-c.

Osorio-Forero, A., Foustoukos, G., Cardis, R., Cherrad, N., Devenoges, C., Fernandez, L. M. J., et al. (2023). Locus coeruleus activity fluctuations set a non-reducible timeframe for mammalian NREM-REM sleep cycles. 2023.05.20.541586. doi: 10.1101/2023.05.20.541586.

Palagini, L., Baglioni, C., Ciapparelli, A., Gemignani, A., and Riemann, D. (2013). REM sleep dysregulation in depression: state of the art. Sleep Med. Rev. 17, 377–390. doi: 10.1016/j.smrv.2012.11.001.

Panula, P., Yang, H. Y., and Costa, E. (1984). Histamine-containing neurons in the rat hypothalamus. Proc. Natl. Acad. Sci. U. S. A. 81, 2572–2576. doi: 10.1073/pnas.81.8.2572.

Rechtschaffen, A., Bergmann, B. M., Gilliland, M. A., and Bauer, K. (1999). Effects of method, duration, and sleep stage on rebounds from sleep deprivation in the rat. Sleep 22, 11–31. doi: 10.1093/sleep/22.1.11.

Ross, R. J., Ball, W. A., Sullivan, K. A., and Caroff, S. N. (1989). Sleep disturbance as the hallmark of posttraumatic stress disorder. Am. J. Psychiatry 146, 697–707. doi: 10.1176/ajp.146.6.697.

Saito, Y. C., Maejima, T., Nishitani, M., Hasegawa, E., Yanagawa, Y., Mieda, M., et al. (2018). Monoamines Inhibit GABAergic Neurons in Ventrolateral Preoptic Area That Make Direct Synaptic Connections to Hypothalamic Arousal Neurons. J. Neurosci. Off. J. Soc. Neurosci. 38, 6366–6378. doi: 10.1523/JNEUROSCI.2835-17.2018.

Sallanon, M., Denoyer, M., Kitahama, K., Aubert, C., Gay, N., and Jouvet, M. (1989). Long-lasting insomnia induced by preoptic neuron lesions and its transient reversal by muscimol injection into the posterior hypothalamus in the cat. Neuroscience 32, 669–683. doi: 10.1016/0306-4522(89)90289-3.

Schott, A. L., Baik, J., Chung, S., and Weber, F. (2023). A medullary hub for controlling REM sleep and pontine waves. Nat. Commun. 14, 3922. doi: 10.1038/s41467-023-39496-0.

Shea, J. L., Mochizuki, T., Sagvaag, V., Aspevik, T., Bjorkum, A. A., and Datta, S. (2008). Rapid eye movement (REM) sleep homeostatic regulatory processes in the rat: changes in the sleep-wake stages and electroencephalographic power spectra. Brain Res. 1213, 48–56. doi: 10.1016/j.brainres.2008.03.062.

Sherin, J. E., Shiromani, P. J., McCarley, R. W., and Saper, C. B. (1996). Activation of Ventrolateral Preoptic Neurons During Sleep. Science 271, 216–219. doi: 10.1126/science.271.5246.216.

Siegel, J., and Gordon, T. P. (1965). PARODOXICAL SLEEP: DEPRIVATION IN THE CAT. Science 148, 978–980. doi: 10.1126/science.148.3672.978.

Smith, J., Honig-Frand, A., Antila, H., Choi, A., Kim, H., Beier, K. T., et al. (2023). Regulation of stress-induced sleep fragmentation by preoptic glutamatergic neurons. Curr. Biol. 0. doi: 10.1016/j.cub.2023.11.035.

Steininger, T. L., Alam, Md. N., Gong, H., Szymusiak, R., and McGinty, D. (1999). Sleep–waking discharge of neurons in the posterior lateral hypothalamus of the albino rat. Brain Res. 840, 138–147. doi: 10.1016/S0006-8993(99)01648-0.

Steininger, T. L., Gong, H., McGinty, D., and Szymusiak, R. (2001). Subregional organization of preoptic area/anterior hypothalamic projections to arousal-related monoaminergic cell groups. J. Comp. Neurol. 429, 638–653.

Stucynski, J. A., Schott, A. L., Baik, J., Chung, S., and Weber, F. (2022). Regulation of REM sleep by inhibitory neurons in the dorsomedial medulla. Curr. Biol. 32, 37–50.e6. doi: 10.1016/j.cub.2021.10.030.

Szymusiak, R., Alam, N., Steininger, T. L., and McGinty, D. (1998). Sleep-waking discharge patterns of ventrolateral preoptic/anterior hypothalamic neurons in rats. Brain Res. 803, 178–188. doi: 10.1016/s0006-8993(98)00631-3.

Takahashi, K., Lin, J.-S., and Sakai, K. (2006). Neuronal Activity of Histaminergic Tuberomammillary Neurons During Wake–Sleep States in the Mouse. J. Neurosci. 26, 10292–10298. doi: 10.1523/JNEUROSCI.2341-06.2006.

Takahashi, K., Lin, J.-S., and Sakai, K. (2009). Characterization and mapping of sleep-waking specific neurons in the basal forebrain and preoptic hypothalamus in mice. Neuroscience 161, 269–292. doi: 10.1016/j.neuroscience.2009.02.075.

Taniguchi, H., He, M., Wu, P., Kim, S., Paik, R., Sugino, K., et al. (2011). A resource of Cre driver lines for genetic targeting of GABAergic neurons in cerebral cortex. Neuron 71, 995–1013. doi: 10.1016/j.neuron.2011.07.026.

Tervo, D. G. R., Hwang, B.-Y., Viswanathan, S., Gaj, T., Lavzin, M., Ritola, K. D., et al. (2016). A Designer AAV Variant Permits Efficient Retrograde Access to Projection Neurons. Neuron 92, 372–382. doi: 10.1016/j.neuron.2016.09.021.

Torontali, Z. A., Fraigne, J. J., Sanghera, P., Horner, R., and Peever, J. (2019). The Sublaterodorsal Tegmental Nucleus Functions to Couple Brain State and Motor Activity during REM Sleep and Wakefulness. Curr. Biol. CB 29, 3803–3813.e5. doi: 10.1016/j.cub.2019.09.026.

Uygun, D. S., Ye, Z., Zecharia, A. Y., Harding, E. C., Yu, X., Yustos, R., et al. (2016). Bottom-Up versus Top-Down Induction of Sleep by Zolpidem Acting on Histaminergic and Neocortex Neurons. J. Neurosci. 36, 11171–11184. doi: 10.1523/JNEUROSCI.3714-15.2016.

Van Dort, C. J., Zachs, D. P., Kenny, J. D., Zheng, S., Goldblum, R. R., Gelwan, N. A., et al. (2015). Optogenetic activation of cholinergic neurons in the PPT or LDT induces REM sleep. Proc. Natl. Acad. Sci. U. S. A. 112, 584–589. doi: 10.1073/pnas.1423136112.

Vanni-Mercier, G., Gigout, S., Debilly, G., and Lin, J. S. (2003). Waking selective neurons in the posterior hypothalamus and their response to histamine H3-receptor ligands: an electrophysiological study in freely moving cats. Behav. Brain Res. 144, 227–241. doi: 10.1016/s0166-4328(03)00091-3.

Verret, L., Fort, P., Gervasoni, D., Léger, L., and Luppi, P.-H. (2006). Localization of the neurons active during paradoxical (REM) sleep and projecting to the locus coeruleus noradrenergic neurons in the rat. J. Comp. Neurol. 495, 573–586. doi: 10.1002/cne.20891.

Von Economo, C. (1930). Sleep as a problem of localization. J. Nerv. Ment. Dis. 71, 249–259. doi: 10.1097/00005053-193003000-00007.

Watanabe, T., Taguchi, Y., Hayashi, H., Tanaka, J., Shiosaka, S., Tohyama, M., et al. (1983). Evidence for the presence of a histaminergic neuron system in the rat brain: an immunohistochemical analysis. Neurosci. Lett. 39, 249–254. doi: 10.1016/0304-3940(83)90308-7.

Weber, F., Machens, C. K., and Borst, A. (2010). Spatiotemporal response properties of optic-flow processing neurons. Neuron 67, 629–642. doi: 10.1016/j.neuron.2010.07.017.

Weber, F., Chung, S., Beier, K. T., Xu, M., Luo, L., and Dan, Y. (2015). Control of REM sleep by ventral medulla GABAergic neurons. Nature 526, 435–438. doi: 10.1038/nature14979.

Weber, F., Hoang Do, J. P., Chung, S., Beier, K. T., Bikov, M., Saffari Doost, M., et al. (2018). Regulation of REM and Non-REM Sleep by Periaqueductal GABAergic Neurons. Nat. Commun. 9, 354. doi: 10.1038/s41467-017-02765-w.

Wiegert, J. S., Mahn, M., Prigge, M., Printz, Y., and Yizhar, O. (2017). Silencing Neurons: Tools, Applications, and Experimental Constraints. Neuron 95, 504–529. doi: 10.1016/j.neuron.2017.06.050.

Williams, R. H., Chee, M. J. S., Kroeger, D., Ferrari, L. L., Maratos-Flier, E., Scammell, T. E., et al. (2014). Optogenetic-mediated release of histamine reveals distal and autoregulatory mechanisms for controlling arousal. J. Neurosci. Off. J. Soc. Neurosci. 34, 6023–6029. doi: 10.1523/JNEUROSCI.4838-13.2014.

Zecharia, A. Y., Yu, X., Götz, T., Ye, Z., Carr, D. R., Wulff, P., et al. (2012). GABAergic Inhibition of Histaminergic Neurons Regulates Active Waking But Not the Sleep–Wake Switch or Propofol-Induced Loss of Consciousness. J. Neurosci. 32, 13062–13075. doi: 10.1523/JNEUROSCI.2931-12.2012.

Zhang, Y., Rózsa, M., Liang, Y., Bushey, D., Wei, Z., Zheng, J., et al. (2023). Fast and sensitive GCaMP calcium indicators for imaging neural populations. Nature 615, 884–891. doi: 10.1038/s41586-023-05828-9.

Zhang, Z., Ferretti, V., Güntan, İ., Moro, A., Steinberg, E. A., Ye, Z., et al. (2015). Neuronal ensembles sufficient for recovery sleep and the sedative actions of α2 adrenergic agonists. Nat. Neurosci. 18, 553–561. doi: 10.1038/nn.395

Zhou, Y., Lai, C. S. W., Bai, Y., Li, W., Zhao, R., Yang, G., et al. (2020). REM sleep promotes experience-dependent dendritic spine elimination in the mouse cortex. Nat. Commun. 11, 4819. doi: 10.1038/s41467-020-18592-5.

