## Supplementary material for "Homeostatic regulation of REM sleep by the preoptic area of the hypothalamus": Table S1

**Table S1. Statistical analysis**

| | | | | | | |
|---|---|---|---|---|---|---|
| **Figure 1C - ΔF/F (%)** | | | | | | |
| **One-way RM ANOVA** | | **F** | **DOF** | **p** | **ηp2** | **Sample size** |
| Main effect of brain state | | 19.17 | (2,18) | 0.0009 | 0.6805 | n=10 mice; 24 recordings |
| Bonferroni's multiple comparisons test | **Comparisons** | **Mean Diff.** | **CI (%)** | **p (adj.)** | **T** | |
| | REM – Wake | 9.774 | (3.17, 16.38) | 0.0056 | 4.341 | |
| | REM – NREM | 11.48 | (4.147, 18.82) | 0.0039 | 4.592 | |
| | Wake – NREM | 1.708 | (-0.7317, 4.148) | 0.2106 | 2.054 | |
| **Figure 1C - ΔF/F (z-scored)** | | | | | | |
| **One-way RM ANOVA** | | **F** | **DOF** | **p** | **ηp2** | |
| Main effect of brain state | | 96.76 | (2,18) | 2.00E-06 | 0.9149 | |
| Bonferroni's multiple comparisons test | **Comparisons** | **Mean Diff.** | **CI (%)** | **p (adj.)** | **T** | |
| | REM – Wake | 2.717 | (1.731, 3.703) | 6.00E-05 | 8.081 | |
| | REM – NREM | 3.072 | (2.54, 3.604) | 1.00E-07 | 16.95 | |
| | Wake – NREM | 0.3551 | (-0.1466, 0.8568) | 0.203 | 2.076 | |
| **Figure 1D - ΔF/F NREMs to REMs (z-scored)** | | | | | | |
| **One-way RM ANOVA** | | **F** | **DOF** | **p** | **ηp2** | **Sample size** |
| Main effect of time | | 248.1492 | (8,72) | 3.18E-49 | 0.8695 | n=10 mice; 24 recordings |
| Holm-Bonferroni correction Baseline: -60 to -50s | **Comparisons** | **p (adj.)** | **T** | **BF10** | **Hedges' g** | |
| | Baseline – -50 to -40 s | 0.9382 | -0.7558 | 0.392 | -0.1073 | |
| | Baseline – -40 to -30 s | 0.0419 | -3.0426 | 4.751 | -0.7473 | |
| | Baseline – -30 to -20 s | 0.0032 | -5.5032 | 92.255 | -1.5715 | |
| | Baseline – -20 to -10 s | 0.0001 | -8.8099 | 2041.41 | -2.7052 | |
| | Baseline – -10 to -0 s | 1.38E-05 | -12.0058 | 19410 | -3.6414 | |
| | Baseline – 0 to 10 s | 4.06E-08 | -25.0662 | 5660000 | -6.733 | |
| | Baseline – 10 to 20 s | 4.06E-08 | -25.0011 | 5545000 | -7.3889 | |
| | Baseline – 20 to 30 s | 4.40E-08 | -24.6629 | 4982000 | -6.7371 | |
| **Figure 1D - ΔF/F NREMs to wake** | | | | | | |
| **One-way RM ANOVA** | | **F** | **DOF** | **p** | **ηp2** | **Sample size** |
| Main effect of time | | 8.1168 | (8,72) | 9.50E-08 | 0.4468 | n=10 mice; 24 recordings |
| Holm-Bonferroni correction Baseline: -60 to -50s | **Comparisons** | **p (adj.)** | **T** | **BF10** | **Hedges' g** | |
| | Baseline – -50 to -40 s | 1.0000 | 0.135 | 0.311 | 0.0389 | |
| | Baseline – -40 to -30 s | 1.0000 | -0.4897 | 0.342 | -0.1532 | |
| | Baseline – -30 to -20 s | 0.7045 | -2.3398 | 1.941 | -0.7912 | |
| | Baseline – -20 to -10 s | 0.5552 | -2.5221 | 2.441 | -0.852 | |
| | Baseline – -10 to -0 s | 0.0093 | -5.707 | 114.97 | -2.0572 | |
| | Baseline – 0 to 10 s | 0.0106 | -5.5549 | 97.59 | -2.6474 | |
| | Baseline – 10 to 20 s | 0.0993 | -3.8393 | 13.089 | -1.7162 | |
| | Baseline – 20 to 30 s | 1.0000 | -1.7502 | 0.961 | -0.7732 | |
| **Figure 1E - ΔF/F during NREMs** | | | | | | |
| **Paired t tests** | | **Time** | | **p** | **p (adj.)** | **Sample size** |

| | | | | | |
|---|---|---|---|---|---|
| Bonferroni's correction Baseline: 0 to 20% | 20 - 40% | | 2.04E-09 | 8.14E-09 | 10 mice; 24 recordings; n=566 events |
| | 40 - 60% | | 1.66E-12 | 6.63E-12 | |
| | 60 - 80% | | 1.00E-13 | 4.01E-13 | |
| | 80 - 100% | | 1.34E-38 | 5.36E-38 | |
| **Figure 2C - REMs amount** | | | | | |
| **Mixed-effects ANOVA** | **F** | **DOF** | **p** | **ηp2** | **Sample size** |
| Main effect of virus | 3.902 | (1,19) | 0.0629 | 0.1041 | n=12 SwiChR++; n=9 eYFP; 116 recordings |
| Main effect of laser | 18.87 | (1,19) | 0.0003 | 0.1755 | |
| Laser x virus | 9.388 | (1,19) | 0.0064 | 0.3163 | |

| | **Comparisons** | **Mean Diff.** | **CI (%)** | **p (adj.)** | **T** |
|---|---|---|---|---|---|
| Bonferroni's multiple comparisons test | SwiChR++: With laser – W/o laser | -2.119 | (-3.031, -1.208) | 3.00E-05 | 5.658 |
| | eYFP: With laser – W/o laser | -0.3661 | (-1.419, 0.6865) | 0.8158 | 0.8463 |
| | With laser: SwiChR++ – eYFP | -1.941 | (-3.365, -0.5173) | 0.0058 | 3.182 |
| | W/o laser: SwiChR++ – eYFP | -0.1878 | (-1.611, 1.236) | 1.0000 | 0.3078 |

| | | | | |
|---|---|---|---|---|
| **Figure 2C - NREMs amount** | | | | |
| **Mixed-effects ANOVA** | **F** | **DOF** | **p** | **ηp2** |
| Main effect of virus | 1.804 | (1,19) | 0.195 | 0.0659 |
| Main effect of laser | 1.266 | (1,19) | 0.2746 | 0.0166 |
| Laser x virus | 0.1629 | (1,19) | 0.691 | 0.003 |
| **Figure 2C - Wake amount** | | | | |
| **Mixed-effects ANOVA** | **F** | **DOF** | **p** | **ηp2** |
| Main effect of virus | 2.153 | (1,19) | 0.1586 | 0.0768 |
| Main effect of laser | 2.906 | (1,19) | 0.1046 | 0.0361 |
| Laser x virus | 0.6689 | (1,19) | 0.4236 | 0.0241 |
| **Figure 2D - Norm. REM EEG power** | | | | |
| **Unpaired t tests** | **Mean Diff.** | **CI (%)** | **p** | **T** |
| δ: SwiChR++ – eYFP | 0.1107 | (0.0034, 0.218) | 0.0432 | 2.166 |
| θ: SwiChR++ – eYFP | 0.0845 | (-0.0228, 0.1917) | 0.1156 | 1.649 |
| σ: SwiChR++ – eYFP | 0.1299 | (0.035, 0.2248) | 0.0099 | 2.865 |
| **Figure 2E - Effect of post laser inhibition on brain state amount** | | | | |
| **Unpaired t tests** | **Mean Diff.** | **CI (%)** | **p** | **T** |
| REMs: SwiChR++ – eYFP | 0.9582 | (-0.6436, 2.560) | 0.2258 | 1.252 |
| NREMs: SwiChR++ – eYFP | -0.6776 | (-4.547, 3.191) | 0.7180 | 0.3666 |
| Wake: SwiChR++ – eYFP | -0.5323 | (-5.250, 4.185) | 0.8158 | 0.2362 |

| | | | | | |
|---|---|---|---|---|---|
| **Figure 3C - Total number of Ca2+ peaks** | | | | | |
| **Two-way RM ANOVA** | **F** | **DOF** | **p** | **ηp2** | **Sample size** |
| Main effect of time | 4.958 | (1,7) | 0.0613 | 0.0349 | |
| Main effect of treatment | 19.56 | (1,7) | 0.0031 | 0.1611 | |
| Time x treatment | 9.229 | (1,7) | 0.0189 | 0.0508 | |

| | Comparisons | Mean Diff. | CI (%) | p (adj.) | T | |
|---|---|---|---|---|---|---|
| Bonferroni's multiple comparisons test | Restr – Baseline (ZT6.5-7.5) | 14.25 | (5.576. 22.92) | 0.0033 | 5.973 | n=8 mice; 16 recordings |
| | Restr – Reb | 9.375 | (0.7007, 18.05) | 0.0341 | 3.93 | |
| | Restr – Baseline (ZT7.5-8.5) | 13.38 | (4.701, 22.05) | 0.0049 | 5.606 | |
| | Baseline (ZT6.5-7.5) – Reb | -4.875 | (-13.55, 3.799) | 0.4819 | 2.043 | |
| | Baseline (ZT6.5-7.5) – Baseline (ZT7.5-8.5) | -0.875 | (-9.549. 7.799) | 1.0000 | 0.3668 | |
| | Reb – Baseline (ZT7.5-8.5) | 4 | (-4.674, 12.67) | 0.8251 | 1.677 | |

**Figure 3C - Number of NREM Ca2+ peaks**

| Two-way RM ANOVA | | F | DOF | p | ηp2 | Sample size |
|---|---|---|---|---|---|---|
| Main effect of time | | 6.674 | (1,7) | 0.0363 | 0.0867 | n=8 mice; 16 recordings |
| Main effect of treatment | | 8.358 | (1,7) | 0.0233 | 0.0738 | |
| Time x treatment | | 19.76 | (1,7) | 0.003 | 0.136 | |
| | **Comparisons** | **Mean Diff.** | **CI (%)** | **p (adj.)** | **T** | |
| Bonferroni's multiple comparisons test | Restr – Baseline (ZT6.5-7.5) | 14 | (4.674, 23.33) | 0.0057 | 5.458 | |
| | Restr – Reb | 14.5 | (5.174, 23.83) | 0.0046 | 5.653 | |
| | Restr – Baseline (ZT7.5-8.5) | 12.38 | (3.049. 21.70) | 0.0115 | 4.825 | |
| | Baseline (ZT6.5-7.5) – Reb | 0.5 | (-8.826, 9.826) | 1.000 | 0.1949 | |
| | Baseline (ZT6.5-7.5) – Baseline (ZT7.5-8.5) | -1.625 | (-10.95, 7.701) | 1.000 | 0.6335 | |
| | Reb – Baseline (ZT7.5-8.5) | -2.125 | -11.45, 7.201) | 1.000 | 0.8285 | |

**Figure 3C - Number of REM Ca2+ peaks**

| Two-way RM ANOVA | F | DOF | p | ηp2 | Sample size |
|---|---|---|---|---|---|
| Main effect of time | 1.882 | (1,7) | 0.2124 | 0.0643 | n=8 mice; 16 recordings |
| Main effect of treatment | 19.72 | (1,7) | 0.003 | 0.1504 | |
| Time x treatment | 4.853 | (1,7) | 0.0634 | 0.1177 | |

**Figure 3D - Amplitude of NREM Ca2+ peaks during restriction**

| Unpaired t tests | Mean Diff. | CI (%) | p | T | Sample size |
|---|---|---|---|---|---|
| Restr – Baseline (ZT6.5-7.5) | 0.0274 | (0.0188, 0.0359) | 6.30E-10 | 6.31 | n=295 restr; n=196 baseline peaks |

**Figure 3E - Amplitude of REM Ca2+ peaks during restriction**

| Unpaired t tests | Mean Diff. | CI (%) | p | T | Sample size |
|---|---|---|---|---|---|
| Restr – Baseline (ZT6.5-7.5) | 0.0424 | (0.0012, 0.0836) | 0.0437 | 2.048 | n=44 restr; n=42 baseline peaks |

**Figure 3F - Amplitude of NREM Ca2+ peaks during rebound**

| Unpaired t tests | Mean Diff. | CI (%) | p | T | Sample size |
|---|---|---|---|---|---|
| Reb– Baseline (ZT7.5-8.5) | 0.0120 | (0.0057, 0.0183) | 0.0002 | 3.744 | n=180 reb; n=196 baseline peaks |
| **Figure 3G - Amplitude of REM Ca2+ peaks during rebound** | | | | | |
| **Unpaired t tests** | **Mean Diff.** | **CI (%)** | **p** | **T** | **Sample size** |
| Reb– Baseline (ZT7.5-8.5) | 0.0127 | (-0.0149, 0.0404) | 0.3641 | 0.9112 | n=84 reb; n=36 baseline peaks |
| **Figure 4C - Effect of laser inhibition during REM restriction (ZT4.5 - 7.5) on brain state amount** | | | | | |
| **Unpaired t tests** | **Mean Diff.** | **CI (%)** | **p** | **T** | **Sample size** |
| REM: SwiChR++ – eYFP | -2.325 | (-4.329, -0.3211) | 0.026 | 2.488 | n=9 SwiChR++; n=7 eYFP |
| NREM: SwiChR++ – eYFP | -2.996 | (-13.30, 7.312) | 0.5431 | 0.6233 | |
| Wake: SwiChR++ – eYFP | 4.17 | (-7.701, 16.04) | 0.4637 | 0.7533 | |
| **Figure 4D - Effect of laser inhibition during REM restriction (ZT4.5 - 7.5) on REM frequency** | | | | | |
| **Unpaired t tests** | **Mean Diff.** | **CI (%)** | **p** | **T** | **Sample size** |
| SwiChR++ – eYFP | -5.487 | (-11.77, 0.7955,) | 0.0821 | 1.873 | n=9 SwiChR++; n=7 eYFP |
| **Figure 4E - Norm. EEG delta, theta and sigma power during REMs restriction (ZT4.5 - 7.5) with laser inhibition** | | | | | |
| **Unpaired t tests** | **Mean Diff.** | **CI (%)** | **p** | **T** | **Sample size** |
| REM: δ SwiChR++ – eYFP | -0.3336 | (-0.5677, -0.0996) | 0.0091 | 3.106 | n=8 SwiChR++; n=6 eYFP |
| REM: θ SwiChR++ – eYFP | -0.1065 | (-0.3035, 0.0906) | 0.2619 | 1.177 | |
| REM: σ SwiChR++ – eYFP | -0.08642 | (-0.3132, 0.1404) | 0.4226 | 0.8302 | |
| NREM: δ SwiChR++ – eYFP | -0.0800 | (-0.2313, 0.0712) | 0.2736 | 1.1430 | n=9 SwiChR++; n=6 eYFP |
| NREM: θ SwiChR++ – eYFP | -0.1614 | (-0.3078, -0.0150) | 0.0332 | 2.381 | |
| NREM: σ SwiChR++ – eYFP | -0.1659 | (-0.321, -0.0107) | 0.038 | 2.31 | |
| **Figure 4F - Brain state amount during REM rebound (ZT7.5 - 8.5)** | | | | | |
| **Unpaired t tests** | **Mean Diff.** | **CI (%)** | **p** | **T** | **Sample size** |
| REM: SwiChR++ – eYFP | -5.082 | (-9.256, -0.9075) | 0.0205 | 2.611 | n=9 SwiChR++; n=7 eYFP |
| NREM: SwiChR++ – eYFP | -1.414 | (-9.097, 6.269) | 0.6989 | 0.3948 | |
| Wake: SwiChR++ – eYFP | 6.195 | (-3.918, 16.31) | 0.21 | 1.314 | |
| **Figure 4G - REM frequency during REM rebound (ZT7.5 - 8.5)** | | | | | |

| Unpaired t tests | Mean Diff. | CI (%) | p | T | Sample size |
|---|---|---|---|---|---|
| SwiChR++ – eYFP | -1.238 | (-4.399, 1.923) | 0.4149 | 0.8401 | n=9 SwiChR++; n=7 eYFP |

**Figure 4H - Norm. EEG delta, theta and sigma power during REM rebound (ZT7.5 - 8.5)**

| Unpaired t tests | Mean Diff. | CI (%) | p | T | Sample size |
|---|---|---|---|---|---|
| REM: δ SwiChR++ – eYFP | -0.1165 | (0.3808, -0.1479) | 0.3607 | 0.9451 | n=8 SwiChR++; n=7 eYFP |
| REM: θ SwiChR++ – eYFP | -0.1751 | (-0.3922, 0.0419) | 0.1073 | 1.695 | |
| REM: σ SwiChR++ – eYFP | -0.0584 | (-0.2629, 0.1462) | 0.5563 | 0.5995 | |
| NREM: δ SwiChR++ – eYFP | -0.0349 | (-0.2290, 0.1593,) | 0.706 | 0.3850 | n=9 SwiChR++; n=7 eYFP |
| NREM: θ SwiChR++ – eYFP | -0.0806 | (-0.2628, 0.1016,) | 0.3587 | 0.9489 | |
| NREM: σ SwiChR++ – eYFP | -0.07111 | (-0.2736, 0.1314,) | 0.4638 | 0.7532 | |

**Figure S1C - ΔF/F (%)**

| One-way RM ANOVA | | F | DOF | p | ηp2 | Sample size |
|---|---|---|---|---|---|---|
| Main effect of brain state | | 6.517 | (2,22) | 0.0162 | 0.372 | n=12 mice; 107 recordings |
| Bonferroni's multiple comparisons test | Comparisons | Mean Diff. | CI (%) | p (adj.) | T | |
| | REM – Wake | 3.011 | (0.4927, 5.530) | 0.0187 | 3.372 | |
| | REM – NREM | 3.225 | (0.4341, 6.883) | 0.0908 | 2.485 | |
| | Wake – NREM | 0.2132 | (-1.806, 2.233) | 1.00 | 0.2977 | |

**Figure S1C - ΔF/F (z-scored)**

| One-way RM ANOVA | | F | DOF | p | ηp2 | Sample size |
|---|---|---|---|---|---|---|
| Main effect of brain state | | 12.31 | (2,22) | 0.0029 | 0.5281 | n=12 mice; 107 recordings |
| Bonferroni's multiple comparisons test | Comparisons | Mean Diff. | CI (%) | p (adj.) | T | |
| | REM – Wake | 1.197 | (0.4864, 1.908) | 0.0018 | 4.75 | |
| | REM – NREM | 1.249 | (0.1691, 2.329) | 0.0227 | 3.262 | |
| | Wake – NREM | 0.05196 | (-0.4622, 0.5662) | 1.00 | 0.285 | |

**Figure S1D - ΔF/F NREMs to REMs (z-scored)**

| One-way RM ANOVA | | F | DOF | p | ηp2 | Sample size |
|---|---|---|---|---|---|---|
| Main effect of time | | 7.160 | (8, 136) | 6.89E-08 | 0.2254 | n=12 mice; 107 recordings |
| Holm-Bonferroni correction Baseline: -60 to -50s | Comparisons | p (adj.) | T | BF10 | Hedges' g | |
| | Baseline – -50 to -40 s | 0.9687 | -1.6228 | 0.732 | -0.2863 | |
| | Baseline – -40 to -30 s | 0.3266 | -2.5938 | 3.13 | -0.6534 | |
| | Baseline – -30 to -20 s | 0.0149 | -3.9141 | 33.98 | -1.1513 | |
| | Baseline – -20 to -10 s | 0.0067 | -4.4643 | 95.561 | -1.5424 | |
| | Baseline – -10 to -0s | 0.0994 | -4.0978 | 47.973 | -1.5385 | |
| | Baseline – 0 to 10 s | 0.0002 | -3.8919 | 32.594 | -1.4571 | |
| | Baseline – 10 to 20 s | 0.0516 | -3.0664 | 7.124 | -1.1493 | |
| | Baseline – 20 to 30s | 0.0361 | -2.2091 | 1.679 | -0.7868 | |

| Figure S1D - ΔF/F NREMs to wake (z-scored) | | | | | | |
|---|---|---|---|---|---|---|
| **One-way RM ANOVA** | | **F** | **DOF** | **p** | **ηp2** | **Sample size** |
| Main effect of time | | 9.9795 | (8, 136) | 7.15E-11 | 0.3137 | n=12 mice; 107 recordings |
| Holm-Bonferroni correction Baseline: -60 to -50s | **Comparisons** | **p (adj.)** | **T** | **BF10** | **Hedges' g** | |
| | Baseline – -50 to -40 s | 1.0000 | 0.0929 | 0.244 | 0.0109 | |
| | Baseline – -40 to -30 s | 1.0000 | -1.0101 | 0.38 | -0.1782 | |
| | Baseline – -30 to -20 s | 0.3133 | -2.7081 | 3.8 | -0.5109 | |
| | Baseline – -20 to -10 s | 0.1752 | -3.0835 | 7.346 | -0.6129 | |
| | Baseline – -10 to -0s | 0.8944 | -1.7431 | 0.855 | -0.4372 | |
| | Baseline – 0 to 10 s | 0.0050 | 4.8473 | 195.709 | 1.7266 | |
| | Baseline – 10 to 20 s | 0.6455 | 2.0935 | 1.408 | 0.7209 | |
| | Baseline – 20 to 30s | 0.5446 | 2.2762 | 1.865 | 0.7958 | |

| Figure S1D - ΔF/F REMs to wake (z-scored) | | | | | | |
|---|---|---|---|---|---|---|
| **One-way RM ANOVA** | | **F** | **DOF** | **p** | **ηp2** | **Sample size** |
| Main effect of time | | 39.1134 | (8, 136) | 1.07E-31 | 0.3293 | n=12 mice; 107 recordings |
| Holm-Bonferroni correction Baseline: -60 to -50s | **Comparisons** | **p (adj.)** | **T** | **BF10** | **Hedges' g** | |
| | Baseline – -50 to -40 s | 0.5099 | -2.1932 | 1.638 | -0.1471 | |
| | Baseline – -40 to -30 s | 0.0730 | -3.2648 | 10.196 | -0.3094 | |
| | Baseline – -30 to -20 s | 0.4171 | -2.3346 | 2.046 | -0.2224 | |
| | Baseline – -20 to -10 s | 0.5099 | -2.1842 | 1.616 | -0.2956 | |
| | Baseline – -10 to -0s | 0.5930 | -2.0212 | 1.264 | -0.3061 | |
| | Baseline – 0 to 10 s | 1.99E-06 | 7.0400 | 9547.365 | 1.0485 | |
| | Baseline – 10 to 20 s | 2.06E-08 | 9.8071 | 629000 | 2.0176 | |
| | Baseline – 20 to 30s | 8.33E-07 | 7.5238 | 21070 | 1.7406 | |

| Figure S1D - ΔF/Fwake to NREMs (z-scored) | | | | | | |
|---|---|---|---|---|---|---|
| **One-way RM ANOVA** | | **F** | **DOF** | **p** | **ηp2** | **Sample size** |
| Main effect of time | | 13.17495741 | (8, 136) | 5.71E-14 | 0.2193 | n=12 mice; 107 recordings |
| Holm-Bonferroni correction Baseline: -60 to -50s | **Comparisons** | **p (adj.)** | **T** | **BF10** | **Hedges' g** | |
| | Baseline – -50 to -40 s | 1.0000 | 0.9997 | 0.376 | 0.0596 | |
| | Baseline – -40 to -30 s | 1.0000 | 1.5243 | 0.648 | 0.1070 | |
| | Baseline – -30 to -20 s | 0.4493 | 2.2974 | 1.928 | 0.2584 | |
| | Baseline – -20 to -10 s | 0.0010 | 5.5047 | 656.926 | 0.6493 | |
| | Baseline – -10 to -0s | 9.10E-09 | 13.1201 | 36970000 | 1.5180 | |
| | Baseline – 0 to 10 s | 0.0001 | 7.1061 | 10650 | 1.7551 | |
| | Baseline – 10 to 20 s | 0.0054 | 4.6368 | 132.06 | 1.3416 | |
| | Baseline – 20 to 30s | 0.2818 | 2.6283 | 3.318 | 0.9186 | |

| Figure S2C - ΔF/F (%) | | | | | | |
|---|---|---|---|---|---|---|
| **One-way RM ANOVA** | | **F** | **DOF** | **p** | **ηp2** | **Sample size** |
| Main effect of brain state | | 19.68 | (2,20) | 5.23E-04 | 0.6631 | |
| Bonferroni's multiple comparisons test | **Comparisons** | **Mean Diff.** | **CI (%)** | **p (adj.)** | **T** | |
| | REM – Wake | -3.557 | (-5.835, -1.28) | 0.0035 | 4.482 | |
| | REM – NREM | -0.3531 | (-1.222, 0.516) | 0.8121 | 1.166 | |

| | | | | | | |
|---|---|---|---|---|---|---|
| | Wake – NREM | 3.204 | (1.280, 5.129) | 0.0022 | 4.779 | |
| **Figure S2C - ΔF/F (z-scored)** | | | | | | n=11 mice; 41 recordings |
| **One-way RM ANOVA** | | **F** | **DOF** | **p** | **ηp2** | |
| Main effect of brain state | | 147.2 | (2,20) | 1.862E-11 | 0.9364 | |
| Bonferroni's multiple comparisons test | **Comparisons** | **Mean Diff.** | **CI (%)** | **p (adj.)** | **T** | |
| | REM – Wake | -1.514 | (-1.837, -1.192) | 2.95E-07 | 13.46 | |
| | REM – NREM | -0.1439 | (-0.4172, 0.1295) | 0.49 | 1.511 | |
| | Wake – NREM | 1.37 | (1.134, 1.607) | 3.81E-08 | 16.66 | |
| **Figure S2D - ΔF/F NREMs to REMs (z-scored)** | | | | | | |
| **One-way RM ANOVA** | | **F** | **DOF** | **p** | **ηp2** | **Sample size** |
| Main effect of time | | 14.1919 | (8,80) | 1.18E-12 | 0.4003 | n=11 mice; 41 recordings |
| Holm-Bonferroni correction Baseline: -60 to -50s | **Comparisons** | **p (adj.)** | **T** | **BF10** | **Hedges' g** | |
| | Baseline – -50 to -40 s | 1.0000 | 1.3200 | 0.597 | 0.3013 | |
| | Baseline – -40 to -30 s | 0.8771 | 2.0335 | 1.334 | 0.4797 | |
| | Baseline – -30 to -20 s | 0.7413 | 2.3087 | 1.895 | 0.5665 | |
| | Baseline – -20 to -10 s | 1.0000 | 1.5964 | 0.797 | 0.3430 | |
| | Baseline – -10 to -0 s | 1.0000 | 0.0139 | 0.298 | 0.0034 | |
| | Baseline – 0 to 10 s | 1.0000 | -1.3712 | 0.628 | -0.5128 | |
| | Baseline – 10 to 20 s | 0.3201 | -2.9223 | 4.316 | -1.1692 | |
| | Baseline – 20 to 30 s | 0.1247 | -3.5060 | 9.596 | -1.5181 | |
| **Figure S2D - ΔF/F NREMs to wake (z-scored)** | | | | | | |
| **One-way RM ANOVA** | | **F** | **DOF** | **p** | **ηp2** | **Sample size** |
| Main effect of time | | 111.2735 | (8,80) | 4.28E-40 | 0.8335 | n=11 mice; 41 recordings |
| Holm-Bonferroni correction Baseline: -60 to -50s | **Comparisons** | **p (adj.)** | **T** | **BF10** | **Hedges' g** | |
| | Baseline – -50 to -40 s | 1.0000 | 1.1771 | 0.523 | 0.1151 | |
| | Baseline – -40 to -30 s | 0.1716 | 3.0063 | 4.841 | 0.4314 | |
| | Baseline – -30 to -20 s | 0.0643 | 3.7499 | 13.37 | 0.5532 | |
| | Baseline – -20 to -10 s | 0.5568 | 2.2209 | 1.692 | 0.4427 | |
| | Baseline – -10 to -0 s | 0.2252 | 2.8008 | 3.658 | 0.3629 | |
| | Baseline – 0 to 10 s | 1.41E-05 | -11.3691 | 30510 | -3.1566 | |
| | Baseline – 10 to 20 s | 1.35E-05 | -11.6411 | 37120 | -4.1001 | |
| | Baseline – 20 to 30 s | 1.44E-05 | -11.1641 | 26250 | -4.0837 | |
| **Figure S2D - ΔF/F REMs to wake (z-scored)** | | | | | | |
| **One-way RM ANOVA** | | **F** | **DOF** | **p** | **ηp2** | **Sample size** |
| Main effect of time | | 32.6242 | (8,80) | 3.69E-22 | 0.5089 | n=11 mice; 41 recordings |
| Holm-Bonferroni correction Baseline: -60 to -50s | **Comparisons** | **p (adj.)** | **T** | **BF10** | **Hedges' g** | |
| | Baseline – -50 to -40 s | 1.0000 | 0.1931 | 0.302 | 0.0203 | |
| | Baseline – -40 to -30 s | 1.0000 | -1.4257 | 0.664 | -0.1436 | |
| | Baseline – -30 to -20 s | 0.6239 | -2.3364 | 1.965 | -0.2845 | |
| | Baseline – -20 to -10 s | 1.0000 | -1.5107 | 0.726 | -0.2829 | |
| | Baseline – -10 to -0s | 1.0000 | -1.0521 | 0.469 | -0.1834 | |
| | Baseline – 0 to 10 s | 0.0008 | -7.4133 | 1034.434 | -2.6524 | |

| | | | | | | |
|---|---|---|---|---|---|---|
| | Baseline – 10 to 20 s | 0.0020 | -6.5109 | 401.011 | -2.0901 | |
| | Baseline – 20 to 30s | 0.0524 | -3.9765 | 18.141 | -1.2891 | |
| **Figure S2D - ΔF/F Wake to NREM (z-scored)** | | | | | | |
| **One-way RM ANOVA** | | **F** | **DOF** | **p** | **ηp2** | **Sample size** |
| Main effect of time | | 35.7154 | (8,80) | 2.38E-23 | 0.4886 | n=11 mice; 41 recordings |
| Holm-Bonferroni correction Baseline: -60 to -50s | **Comparisons** | **p (adj.)** | **T** | **BF10** | **Hedges' g** | |
| | Baseline – -50 to -40 s | 0.0722 | -3.5187 | 9.763 | -0.5254 | |
| | Baseline – -40 to -30 s | 0.0743 | -3.4133 | 8.455 | -0.8971 | |
| | Baseline – -30 to -20 s | 0.0087 | -5.1053 | 77.953 | -1.4173 | |
| | Baseline – -20 to -10 s | 0.0099 | -4.9858 | 67.18 | -1.5718 | |
| | Baseline – -10 to -0s | 0.3056 | -2.3862 | 2.098 | -0.8628 | |
| | Baseline – 0 to 10 s | 9.36E-02 | 3.2084 | 6.386 | 1.0398 | |
| | Baseline – 10 to 20 s | 7.43E-02 | 3.4531 | 8.928 | 1.1135 | |
| | Baseline – 20 to 30s | 3.46E-02 | 4.0542 | 20.125 | 1.1242 | |
| **Figure S2E - ΔF/F during NREMs** | | | | | | |
| **Paired t tests** | | **Time** | | **p** | **p (adj.)** | **Sample size** |
| Bonferroni's correction Baseline: 0 to 20% | | 20 - 40% | | 1.34E-33 | 5.36E-33 | 11 mice; 41 recordings; n=889 events |
| | | 40 - 60% | | 3.11E-46 | 1.24E-45 | |
| | | 60 - 80% | | 4.38E-61 | 1.75E-60 | |
| | | 80 - 100% | | 6.48E-71 | 2.59E-70 | |
| **Figure S3A - REM duration** | | | | | | |
| **Mixed-effects ANOVA** | | **F** | **DOF** | **p** | **ηp2** | **Sample size** |
| Main effect of virus | | 0.2531 | (1,19) | 0.6207 | 0.0084 | n=12 SwiChR++; n=9 eYFP; 116 recordings |
| Main effect of laser | | 0.0040 | (1,19) | 0.9501 | 0.0005 | |
| Laser x virus | | 2.629 | (1,19) | 0.1214 | 0.0831 | |
| **Figure S3A - NREM duration** | | | | | | |
| **Mixed-effects ANOVA** | | **F** | **DOF** | **p** | **ηp2** | |
| Main effect of virus | | 10.34 | (1,19) | 0.0046 | 0.2803 | |
| Main effect of laser | | 0.3348 | (1,19) | 0.5697 | 0.0038 | |
| Laser x virus | | 0.0081 | (1,19) | 0.929 | 0.0002 | |
| **Figure S3A - Wake duration** | | | | | | |
| **Mixed-effects ANOVA** | | **F** | **DOF** | **p** | **ηp2** | |
| Main effect of virus | | 0.192 | (1,19) | 0.6662 | 0.0070 | |
| Main effect of laser | | 2.982 | (1,19) | 0.1004 | 0.0455 | |
| Laser x virus | | 2.399 | (1,19) | 0.1379 | 0.0424 | |
| **Figure S3B - REM frequency** | | | | | | |
| **Mixed-effects ANOVA** | | **F** | **DOF** | **p** | **ηp2** | |
| Main effect of virus | | 2.901 | (1,19) | 0.1048 | 0.0712 | |
| Main effect of laser | | 3.863 | (1,19) | 0.0642 | 0.0858 | |
| Laser x virus | | 1.285 | (1,19) | 0.2711 | 0.0604 | |
| **Figure S3B - NREM frequency** | | | | | | |
| **Mixed-effects ANOVA** | | **F** | **DOF** | **p** | **ηp2** | |

| | | | | | |
|---|---|---|---|---|---|
| Main effect of virus | 3.715 | (1,19) | 0.069 | 0.135 | |
| Main effect of laser | 0.0762 | (1,19) | 0.7854 | 0.001 | |
| Laser x virus | 0.0655 | (1,19) | 0.8008 | 0.008 | |
| **Figure S3B - Wake frequency** | | | | | |
| **Mixed-effects ANOVA** | **F** | **DOF** | **p** | **ηp2** | |
| Main effect of virus | 9.142 | (1,19) | 0.007 | 0.2808 | |
| Main effect of laser | 0.0006 | (1,19) | 0.98 | 0.0078 | |
| Laser x virus | 0.7833 | (1,19) | 0.7833 | 0.0002 | |
| **Figure S3C - REM EEG power** | | | | | |
| **Paired t tests** | **Mean Diff.** | **CI (%)** | **p** | **T** | **Sample size** |
| δ: SwiChR++ with laser - w/o laser | 153.7 | (57.45, 250.1) | 0.0048 | 3.514 | n=12 mice; 62 recordings |
| θ: SwiChR++ with laser - w/o laser | 198 | (-35.78, 431.7) | 0.0892 | 1.864 | |
| σ: SwiChR++ with laser - w/o laser | 68.81 | (15.64, 122,) | 0.0158 | 2.848 | |
| **Figure S3C - NREM EEG power** | | | | | |
| **Paired t tests** | **Mean Diff.** | **CI (%)** | **p** | **T** | |
| δ: SwiChR++ with laser - w/o laser | 174.8 | (-686.2, 1036) | 0.6636 | 0.4469 | |
| θ: SwiChR++ with laser - w/o laser | -113.6 | (-361.5, 134.3) | 0.3348 | 1.009 | |
| σ: SwiChR++ with laser - w/o laser | -146.5 | (-309.9, 17.02) | 0.0743 | 1.972 | |
| **Figure S3C - Wake EEG power** | | | | | |
| **Paired t tests** | **Mean Diff.** | **CI (%)** | **p** | **T** | |
| δ: SwiChR++ with laser - w/o laser | -71.88 | (-177.9, 34.13) | 0.1637 | 1.492 | |
| θ: SwiChR++ with laser - w/o laser | -26.43 | (-193.1, 140.3) | 0.7337 | 0.349 | |
| σ: SwiChR++ with laser - w/o laser | -16.81 | (-52.51, 18.88) | 0.3221 | 1.037 | |
| **Figure S3D - Norm. NREM, wake EEG power** | | | | | |
| **Unpaired t tests** | **Mean Diff.** | **CI (%)** | **p** | **T** | **Sample size** |
| NREM: δ SwiChR++ – eYFP | 0.0300 | (-0.1343, 0.1942) | 0.7067 | 0.382 | n=12 SwiChR++; n=9 eYFP; 116 recordings |
| NREM: θ SwiChR++ – eYFP | -0.0365 | (-0.1338, 0.0608) | 0.4418 | 0.7856 | |
| NREM: σ SwiChR++ – eYFP | -0.0574 | (-0.1672, 0.0524) | 0.2877 | 1.094 | |
| Wake δ: SwiChR++ – eYFP | -0.0094 | (-0.0882, 0.0701) | 0.8076 | 0.2469 | |
| Wake θ: SwiChR++ – eYFP | -0.0108 | (-0.1273, 0.1057) | 0.8479 | 0.1944 | |
| Wake σ: SwiChR++ – eYFP | -0.0064 | (-0.0732, 0.0604) | 0.8423 | 0.2017 | |
| **Figure S3E - Effect of post laser inhibition on brain state duration** | | | | | |
| **Unpaired t tests** | **Mean Diff.** | **CI (%)** | **p** | **T** | **Sample size** |
| REMs: SwiChR++ – eYFP | 3.906 | (-7.921, 15.73) | 0.4978 | 0.6912 | n=12 SwiChR++; n=9 eYFP; 63 recordings |
| NREMs: SwiChR++ – eYFP | -18.65 | (-53.20, 15.91) | 0.2728 | 1.129 | |
| Wake: SwiChR++ – eYFP | -34.67 | (-72.58, 3.241) | 0.0708 | 1.914 | |
| **Figure S3F - Effect of post laser inhibition on brain state frequency** | | | | | |

| Unpaired t tests | Mean Diff. | CI (%) | p | T | Sample size |
|---|---|---|---|---|---|
| REMs: SwiChR++ – eYFP | 0.1972 | (-1.200, 1.594) | 0.7709 | 0.2954 | n=12 SwiChR++; n=9 eYFP; 63 recordings |
| NREMs: SwiChR++ – eYFP | 2.04 | (-1.634, 5.713) | 0.2596 | 1.162 | |
| Wake: SwiChR++ – eYFP | 0.7592 | (-1.002, 2.521) | 0.3784 | 0.902 | |

**Figure S4C - REMs amount**

| Mixed-effects ANOVA | | F | DOF | p | ηp2 | Sample size |
|---|---|---|---|---|---|---|
| Main effect of virus | | 2.323 | (1, 16) | 0.147 | 0.0042 | n=10 SwiChR++; n=8 eYFP; 103 recordings |
| Main effect of laser | | 10.34 | (1, 16) | 0.0054 | 0.5563 | |
| Laser x virus | | 4.706 | (1, 16) | 0.0455 | 0.1493 | |
| Bonferroni's multiple comparisons test | **Comparisons** | **Mean Diff.** | **CI (%)** | **p (adj.)** | **T** | |
| | SwiChR++: With laser – W/o laser | 1.591 | (0.6168, 2.565) | 0.0019 | 4.039 | |
| | eYFP: With laser – W/o laser | 0.3091 | (-0.7797, 1.398) | 0.9855 | 0.7021 | |
| | With laser: SwiChR++ – eYFP | 1.324 | (0.0616, 2.586) | 0.0384 | 2.467 | |
| | W/o laser: SwiChR++ – eYFP | 0.0421 | (-1.220, 1.304) | 1.0000 | 0.0784 | |

**Figure S4C - NREMs amount**

| Mixed-effects ANOVA | F | DOF | p | ηp2 |
|---|---|---|---|---|
| Main effect of virus | 0.6674 | (1, 16) | 0.426 | 0.1170 |
| Main effect of laser | 2.671 | (1, 16) | 0.1217 | 0.4085 |
| Laser x virus | 4.364 | (1, 16) | 0.053 | 0.2518 |

**Figure S4C - Wake amount**

| Mixed-effects ANOVA | | F | DOF | p | ηp2 |
|---|---|---|---|---|---|
| Main effect of virus | | 0.2586 | (1, 16) | 0.618 | 0.1053 |
| Main effect of laser | | 3.237 | (1, 16) | 0.0909 | 0.4621 |
| Laser x virus | | 5.126 | (1, 16) | 0.0378 | 0.2591 |
| Bonferroni's multiple comparisons test | **Comparisons** | **Mean Diff.** | **CI (%)** | **p (adj.)** | **T** |
| | SwiChR++: With laser – W/o laser | -8.731 | (-15.82, -1.646) | 0.0154 | 3.047 |
| | eYFP: With laser – W/o laser | 0.9992 | (-6.922, 8.921) | 1.0000 | 0.3119 |
| | With laser: SwiChR++ – eYFP | -3.224 | (-12.34, 5.895) | 0.8237 | 0.8315 |
| | W/o laser: SwiChR++ – eYFP | 6.506 | (-2.613, 15.63) | 0.2062 | 1.678 |

**Figure S4D - Norm. REM, NREM, wake EEG power**

| Unpaired t tests | Mean Diff. | CI (%) | p | T |
|---|---|---|---|---|
| REM: δ SwiChR++ – eYFP | -0.1189 | (-0.2376, -1.01E-04) | 0.0498 | 2.122 |
| REM: θ SwiChR++ – eYFP | -0.0618 | (-0.1705, 0.047) | 0.2461 | 1.204 |

| | | | | | | |
|---|---|---|---|---|---|---|
| REM:<br>σ SwiChR++ – eYFP | | -0.08868 | (-0.1942, 0.0169) | 0.0939 | 1.781 | |
| NREM δ:<br>SwiChR++ – eYFP | | -0.1160 | (-0.2032, -0.0288) | 0.0123 | 2.8200 | |
| NREM θ:<br>SwiChR++ – eYFP | | -0.0926 | ( -0.184, -0.0012) | 0.0474 | 2.148 | |
| NREM σ:<br>SwiChR++ – eYFP | | -0.0617 | (-0.1493, 0.0259) | 0.1551 | 1.492 | |
| Wake δ:<br>SwiChR++ – eYFP | | 0.0562 | (-0.06367, 0.1760) | 0.3352 | 0.9936 | |
| Wake θ:<br>SwiChR++ – eYFP | | -0.0641 | (-0.1835, 0.0554) | 0.2722 | 1.137 | |
| Wake σ:<br>SwiChR++ – eYFP | | -0.006791 | (-0.1009, 0.0874) | 0.8804 | 0.1529 | |
| **Figure S5B - Motor-on frequency** | | | | | | |
| **One-way RM ANOVA** | | **F** | **DOF** | **p** | **ηp2** | **Sample size** |
| Main effect of time | | 5.835 | (5,45) | 0.0061 | 0.3933 | n=10 mice;<br>10 recordings |
| Bonferroni's multiple comparisons test Baseline: ZT1.5 | **Comparisons** | **Mean Diff.** | **CI (%)** | **p (adj.)** | **T** | |
| | ZT1.5 – ZT2.5 | -6.1 | (-12.54, 0.3353) | 0.0656 | 3.081 | |
| | ZT1.5 – ZT3.5 | -9.1 | (-17.46, -0.7447) | 0.0316 | 3.539 | |
| | ZT1.5 – ZT4.5 | -15.8 | (-29.76, -1.845) | 0.0254 | 3.679 | |
| | ZT1.5 – ZT5.5 | -14.4 | (-27.03, -1.765) | 0.0245 | 3.704 | |
| | ZT1.5 – ZT6.5 | -16.4 | (-27.86, -4.944) | 0.006 | 4.652 | |
| **Figure S5D - Effect of REM restriction on brain state amount** | | | | | | |
| **Paired t tests** | | **Mean Diff.** | **CI (%)** | **p** | **T** | **Sample size** |
| REMs: Restriction - Baseline | | -2.748 | (-3.960, -1.536) | 0.0006 | 5.128 | n=10 mice;<br>20 recordings |
| NREMs: Restriction - Baseline | | 7.06 | (-2.791, 16.91) | 0.1394 | 1.621 | |
| Wake: Restriction - Baseline | | -4.631 | (-15.43, 6.168) | 0.3573 | 0.9701 | |
| **Figure S5E - Effect of REM restriction on brain state duration** | | | | | | |
| **Paired t tests** | | **Mean Diff.** | **CI (%)** | **p** | **T** | |
| REMs: Restriction - Baseline | | -49.16 | (-62.34, -35.99) | 1.44E-05 | 8.441 | |
| NREMs: Restriction - Baseline | | -85.41 | (-159.1, -11.77) | 0.0276 | 2.624 | |
| Wake: Restriction - Baseline | | -153.00 | (-433.4, 127.4) | 0.2482 | 1.235 | |
| **Figure S5F - Effect of REM restriction on brain state frequency** | | | | | | |
| **Paired t tests** | | **Mean Diff.** | **CI (%)** | **p** | **T** | |
| REMs: Restriction - Baseline | | 7.533 | (3.370, 11.70) | 0.0027 | 4.093 | |
| NREMs: Restriction - Baseline | | 8.267 | ( 3.336, 13.20) | 0.0043 | 3.793 | |
| Wake: Restriction - Baseline | | 1.217 | (-0.6769, 3.110) | 0.18 | 1.454 | |
| **Figure S5G - Effect of REM restriction on power spectral density (PSD)** | | | | | | |
| **Paired t tests** | | **Mean Diff.** | **CI (%)** | **p** | **T** | |
| REMs δ: Restriction - Baseline | | 160.2 | (20.21, 300.1) | 0.0293 | 2.589 | |
| REMs θ: Restriction - Baseline | | 82.12 | (-245.9, 410.1) | 0.585 | 0.5664 | |
| REMs σ: Restriction - Baseline | | 90.7 | (-9.430, 190.8) | 0.0707 | 2.049 | |
| NREMs δ: Restriction- Baseline | | 82.57 | (-544.7, 709.8) | 0.7726 | 0.2978 | |

| | | | | | |
|---|---|---|---|---|---|
| NREMs θ: Restriction - Baseline | 155.8 | (-28.85, 340.4) | 0.0887 | 1.909 | |
| NREMs σ: Restriction - Baseline | 123.4 | (-0.4201, 247.3) | 0.0506 | 2.254 | |
| Wake δ: Restriction - Baseline | 193.5 | (-69.98, 457.0) | 0.131 | 1.661 | |
| Wake θ: Restriction - Baseline | -6.578 | (-158.1, 145.0) | 0.9239 | 0.09819 | |
| Wake σ: Restriction - Baseline | 22.51 | (-27.99, 73.01) | 0.3397 | 1.008 | |
| **Figure S5H - Effect of REM restriction on rebound brain state amount** | | | | | |
| **Paired t tests** | **Mean Diff.** | **CI (%)** | **p** | **T** | **Sample size** |
| REMs: Rebound - Baseline | 6.444 | (1.135, 11.75) | 0.0226 | 2.746 | |
| NREMs: Rebound - Baseline | 11.06 | ( -8.583, 30.71) | 0.2346 | 1.274 | |
| Wake: Rebound - Baseline | -17.41 | (-42.20, 7.382) | 0.1466 | 1.589 | |
| **Figure S5I - Effect of REM restriction on rebound brain state duration** | | | | | |
| **Paired t tests** | **Mean Diff.** | **CI (%)** | **p** | **T** | |
| REMs: Rebound - Baseline | 4.038 | (-32.54, 40.61 ) | 0.8084 | 0.2497 | |
| NREMs: Rebound - Baseline | 3.1 | (-1.091, 7.291) | 0.1286 | 1.673 | n=10 mice; 20 recordings |
| Wake: Rebound - Baseline | -323.7 | (-945.0, 297.6) | 0.2687 | 1.179 | |
| **Figure S5J - Effect of REM restriction on rebound brain state frequency** | | | | | |
| **Paired t tests** | **Mean Diff.** | **CI (%)** | **p** | **T** | |
| REMs: Rebound - Baseline | 4.1 | (-6.656, -1.544) | 0.0055 | 3.629 | |
| NREMs: Rebound - Baseline | 3.1 | (-1.091, 7.291) | 0.1286 | 1.673 | |
| Wake: Rebound - Baseline | -1.217 | (-3.110, 0.6769) | 0.18 | 1.454 | |
| **Figure S5K - Effect of REM restriction on rebound PSD** | | | | | |
| **Paired t tests** | **Mean Diff.** | **CI (%)** | **p** | **T** | **Sample size** |
| REMs δ: Rebound - Baseline | 164.5 | (-18.47, 347.5) | 0.0719 | 2.073 | |
| REMs θ: Rebound - Baseline | 178.2 | (-183.8, 540.2) | 0.2892 | 1.135 | n=9 mice; 18 recordings |
| REMs σ: Rebound - Baseline | 24.13 | (-75.24, 123.5) | 0.5908 | 0.56 | |
| NREMs δ: Rebound - Baseline | -96.7 | (-703.7, 897.0) | 0.7908 | 0.2733 | |
| NREMs θ: Rebound - Baseline | 113.8 | (-81.05, 308.6) | 0.2191 | 1.321 | |
| NREMs σ: Rebound - Baseline | 55.53 | (-120.6, 231.6) | 0.4938 | 0.7132 | n=10 mice; 20 recordings |
| Wake δ: Rebound - Baseline | 232.9 | (53.88, 412.0) | 0.0164 | 2.943 | |
| Wake θ: Rebound - Baseline | -89.03 | (-301.1, 123.0) | 0.367 | 0.9499 | |
| Wake σ: Rebound - Baseline | 24.97 | (-58.80, 108.7) | 0.5171 | 0.6742 | |
| **Figure S6A - REM restriction for photometry recordings** | | | | | |
| **Paired t tests** | **Mean Diff.** | **CI (%)** | **p** | **T** | **Sample size** |
| REMs Amount: Restr - Baseline | -5.825 | (-8.810, -2.839) | 0.0024 | 4.614 | |
| REMs Duration: Restr - Baseline | -78.87 | (-115.1, -42.61) | 0.0013 | 5.143 | n=8 mice; 16 recordings |
| REMs Frequency: Restr - Baseline | 1.875 | (-2.158, 5.907) | 0.3079 | 1.099 | |
| **Figure S6B - Effect of REM restriction on REM rebound for photometry recordings** | | | | | |
| **Paired t tests** | **Mean Diff.** | **CI (%)** | **p** | **T** | **Sample size** |
| REMs Amount: Reb - Baseline | 7.344 | (2.788, 11.90) | 0.0066 | 3.812 | |
| REMs Duration: Reb - Baseline | -22.6 | (-81.23, 36.03) | 0.3923 | 0.9116 | n=8 mice; 16 recordings |
| REMs Frequency: Reb - Baseline | 4.5 | (1.894, 7.105) | 0.0047 | 4.084 | |
| **Figure S7A - Effect of laser inhibition during REM restriction on brain state duration** | | | | | |

| Unpaired t tests | Mean Diff. | CI (%) | p | T | Sample size |
|---|---|---|---|---|---|
| REMs Duration: SwiChR++ - eYFP | -5.487 | (-11.77, 0.7955) | 0.0821 | 1.873 | n=9 SwiChR++; n=7 eYFP 16 recordings |
| NREMs Duration: SwiChR++ - eYFP | 22.06 | (-33.55, 77.67) | 0.4092 | 0.8508 | |
| Wake Duration: SwiChR++ - eYFP | 13.85 | (-10.84, 38.54) | 0.249 | 1.203 | |
| **Figure S7B - Effect of laser inhibition during REM restriction on brain state frequecy** | | | | | |
| **Unpaired t tests** | **Mean Diff.** | **CI (%)** | **p** | **T** | |
| NREMs Frequency: SwiChR++ - eYFP | -1.138 | (-9.184, 6.909) | 0.7662 | 0.3032 | |
| Wake Frequnecy: SwiChR++ - eYFP | -0.3439 | (-3.511, 2.823) | 0.8192 | 0.2329 | |
| **Figure S7C - Effect of laser inhibition during REM restriction on norm. EEG wake** | | | | | |
| **Unpaired t tests** | **Mean Diff.** | **CI (%)** | **p** | **T** | |
| Wake δ: SwiChR++ – eYFP | -0.0704 | (-0.2969, 0.1561) | 0.5157 | 0.6669 | |
| Wake θ: SwiChR++ – eYFP | -0.0704 | (-0.2619, 0.1211) | 0.4437 | 0.7883 | |
| Wake σ: SwiChR++ – eYFP | -0.0839 | (-0.2780, 0.1104) | 0.3701 | 0.926 | |
| **Figure S7D - Effect of laser inhibition during REM restriction on brain state duration** | | | | | |
| **Unpaired t tests** | **Mean Diff.** | **CI (%)** | **p** | **T** | **Sample size** |
| REMs Duration: SwiChR++ - eYFP | -20.16 | (-56.07, 15.75) | 0.2484 | 1.204 | n=9 SwiChR++; n=7 eYFP 16 recordings |
| NREMs Duration: SwiChR++ - eYFP | -23.57 | (-107.3, 60.14) | 0.5555 | 0.604 | |
| Wake Duration: SwiChR++ - eYFP | 43.98 | (-81.07, 169) | 0.4632 | 0.7543 | |
| **Figure S7E - Effect of laser inhibition during REM restriction on brain state frequecy** | | | | | |
| **Unpaired t tests** | **Mean Diff.** | **CI (%)** | **p** | **T** | |
| NREMs Frequency: SwiChR++ - eYFP | 1.27 | (-3.456, 5.996) | 0.5736 | 0.5763 | |
| Wake Frequnecy: SwiChR++ - eYFP | 2.778 | (-1.983, 7.538) | 0.2312 | 1.252 | |
| **Figure S7H - Effect of laser inhibition during REM restriction on norm. wake EEG** | | | | | |
| **Unpaired t tests** | **Mean Diff.** | **CI (%)** | **p** | **T** | |
| Wake δ: SwiChR++ – eYFP | 0.1418 | (-0.1039, 0.3875) | 0.2362 | 1.238 | |
| Wake θ: SwiChR++ – eYFP | 0.1937 | (-0.0996, 0.4870) | 0.1786 | 1.416 | |
| Wake σ: SwiChR++ – eYFP | 0.1056 | (-0.1008, 0.3121) | 0.291 | 1.097 | |
