## Supplementary material for "Homeostatic regulation of REM sleep by the preoptic area of the hypothalamus": Supple Figures 1-7

### Figure S1

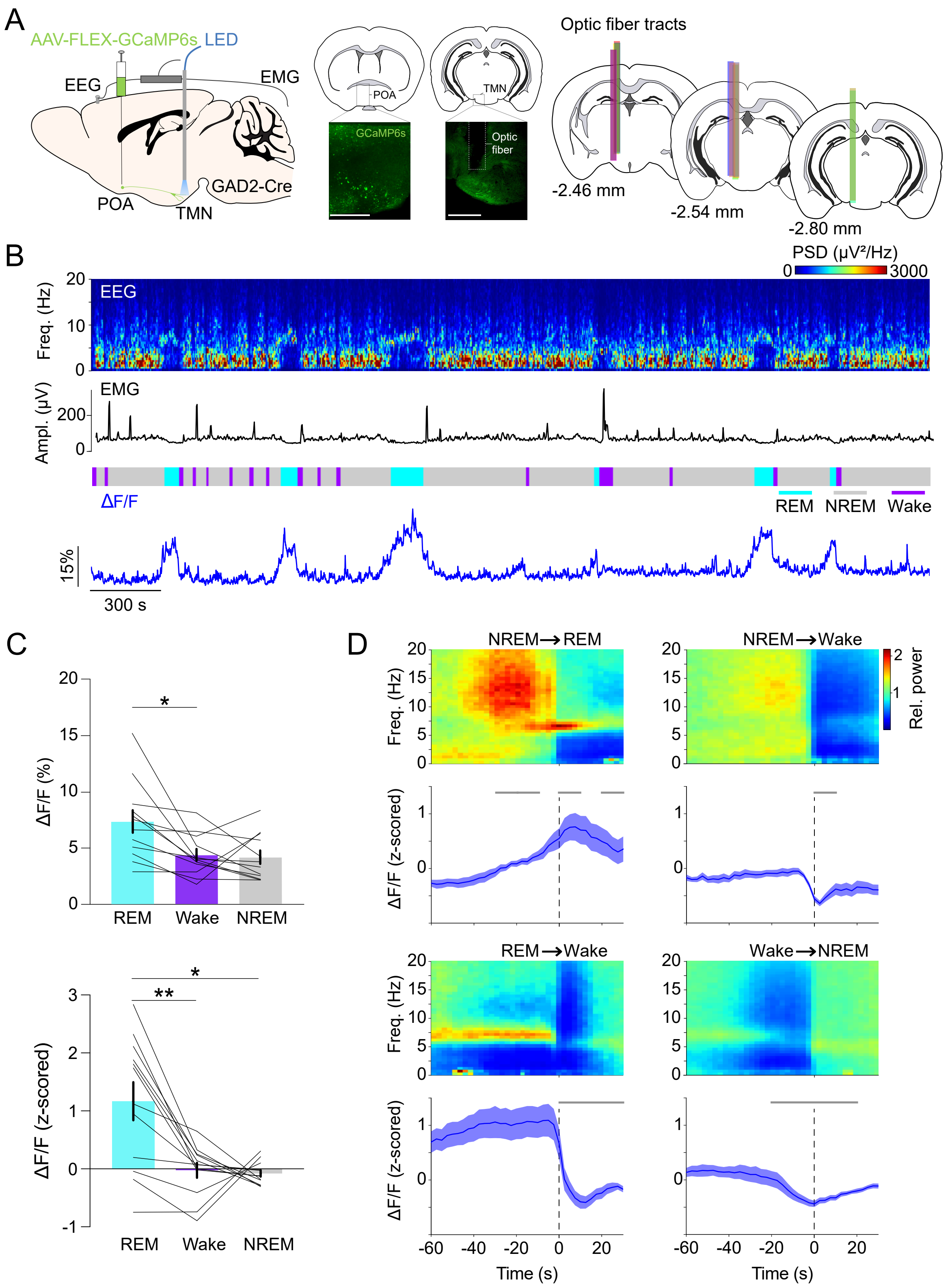

A
AAV-FLEX-GCaMP6s
LED
EEG
EMG
POA
TMN
GAD2-Cre
GCaMP6s
Optic fiber
Optic fiber tracts
-2.46 mm
-2.54 mm
-2.80 mm
B
PSD (μV²/Hz)
0
3000
EEG
Freq. (Hz)
EMG
Ampl. (μV)
ΔF/F
REM
NREM
Wake
15%
300 s
C
ΔF/F (%)
ΔF/F (z-scored)
REM
Wake
NREM
D
NREM→REM
NREM→Wake
REM→Wake
Wake→NREM
Rel. power
Time (s)

**FIGURE S1 | POA$^{GAD2}$→TMN axonal fibers are most active during REMs. Related to Figure 1.**

**(A)** Left, schematic of fiber photometry with simultaneous EEG and EMG recordings. Mouse brain figure adapted from the Allen Reference Atlas - Mouse Brain (atlas.brain-map.org). Middle, fluorescence image of POA and TMN in a GAD2-Cre mouse injected with AAV-FLEX-GCaMP6s into the POA and implanted with an optic fiber in the TMN. Scale bar, 0.5 mm. Right, location of fiber tracts. Each colored bar represents the location of optic fibers for photometry recordings.
**(B)** Example fiber photometry recording. Shown are EEG spectrogram, EMG amplitude, color-coded brain states, and ΔF/F signal.
**(C)** Non-normalized and z-scored ΔF/F activity during REMs, wake, and NREMs. Bars, averages across mice; lines, individual mice; error bars, ± s.e.m. One-way rm ANOVA $p = 0.0162$, 0.0029 for non-normalized ΔF/F and z-scored ΔF/F; pairwise t-tests with Bonferroni correction, non-normalized ΔF/F, $p = 0.0187$ for REMs vs. Wake; z-scored ΔF/F, $p = 0.0018$, 0.0227 for REMs vs. Wake or REMs vs. NREMs.
**(D)** Average EEG spectrogram (top), calcium activity (bottom) during brain state transitions. Shading, ± s.e.m. One-way rm ANOVA, $p = 6.89e-8$ for NREMs→REMs transitions; pairwise t-tests with Holm-Bonferroni correction, $p < 0.0149$ between -30 s and -10 s, $p = 0.0002$ between 0 and 10 s, $p = 0.0361$ between 20 and 30 s. Gray bar, period when ΔF/F activity was significantly different from baseline (-60 to -50 s).

n = 12 mice.
See **Table S1** for the actual p values.

### Figure S2

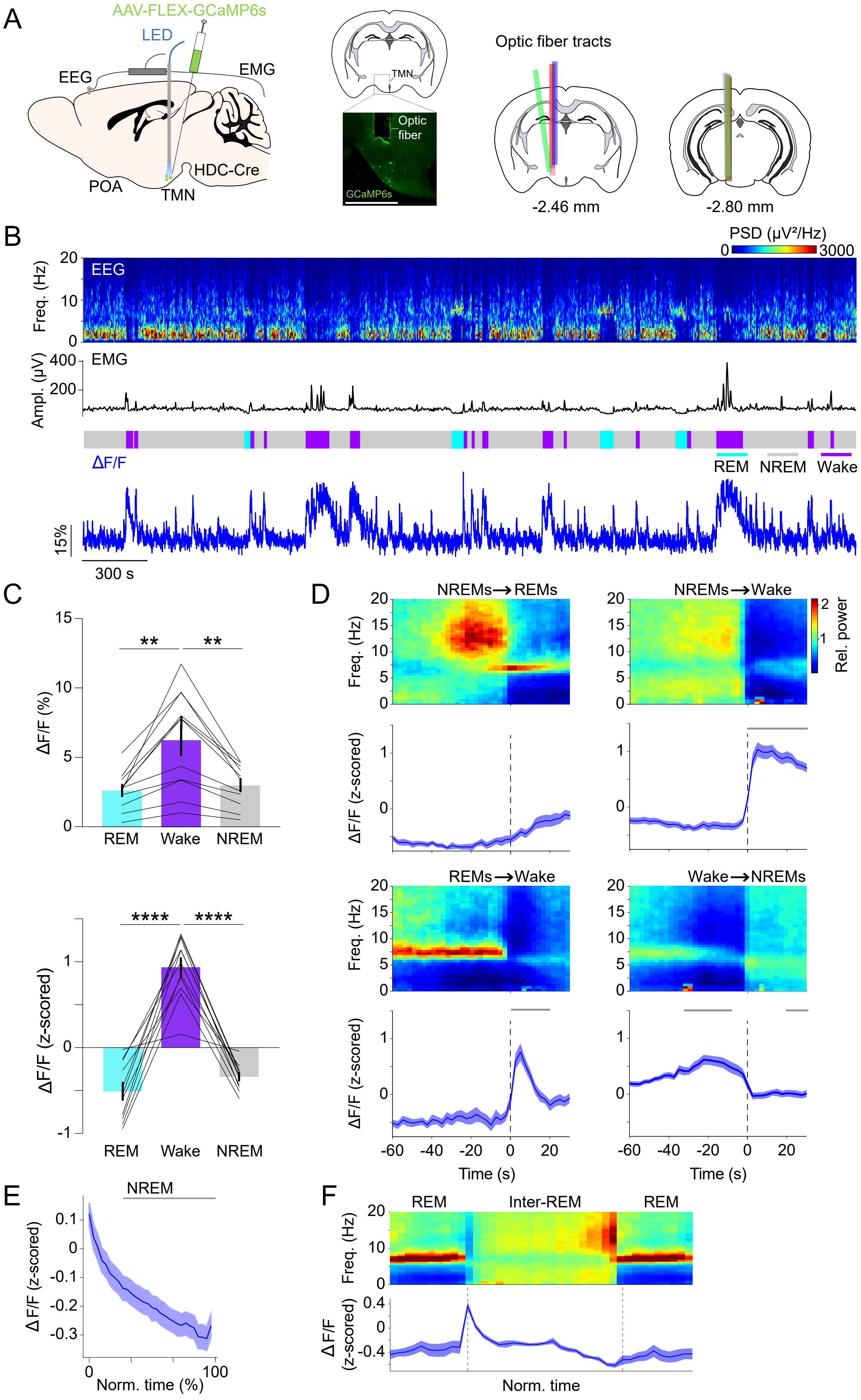
A
AAV-FLEX-GCaMP6s
LED
EEG
EMG
POA
TMN
HDC-Cre
Optic fiber
GCaMP6s
Optic fiber tracts
-2.46 mm
-2.80 mm
B
PSD (µV²/Hz)
0
3000
EEG
Freq. (Hz)
EMG
Ampl. (µV)
ΔF/F
REM
NREM
Wake
15%
300 s
C
ΔF/F (%)
ΔF/F (z-scored)
REM
Wake
NREM
**
****
D
NREMs→REMs
NREMs→Wake
REMs→Wake
Wake→NREMs
Rel. power
Freq. (Hz)
ΔF/F (z-scored)
Time (s)
E
NREM
ΔF/F (z-scored)
Norm. time (%)
F
REM
Inter-REM
REM
Freq. (Hz)
ΔF/F (z-scored)
Norm. time

**FIGURE S2 | TMN$^{HIS}$ neurons are least active during sleep. Related to Figure 1.**

**(A)** Left, schematic of fiber photometry with simultaneous EEG and EMG recordings. Middle, fluorescence image of TMN in a HDC-Cre mouse injected with AAV-FLEX-GCaMP6s and implanted with an optic fiber in the TMN. Scale bar, 1 mm. Right, location of fiber tracts. Each colored bar represents the location of optic fibers for photometry recordings.

**(B)** Example fiber photometry recording. Shown are EEG spectrogram, EMG amplitude, color-coded brain states, and ΔF/F signal.

**(C)** Non-normalized and z-scored ΔF/F activity during REMs, wake, and NREMs. Bars, averages across mice; lines, individual mice; error bars, ± s.e.m. One-way rm ANOVA p = 5.23e-4, 1.862e-11 for non-normalized ΔF/F and z-scored ΔF/F; pairwise t-tests with Bonferroni correction, non-normalized ΔF/F, p = 0.0035, 0.0022 for REMs vs. wake or wake vs. NREMs; z-scored ΔF/F, p = 2.95e-7, 3.81e-8. n = 11 mice.

**(D)** Average EEG spectrogram (top), and calcium activity (bottom) during brain state transitions. Shading, ± s.e.m. One-way rm ANOVA, p = 4.28e-40, 3.69e-22 for NREMs→wake and REMs→wake; pairwise t-tests with Holm-Bonferroni correction, NREMs→wake p < 1.44e-05 between 0 s and 30 s, REMs→wake p < 0.002 between 0 s and 20 s. Gray bar, period when ΔF/F activity was significantly different from baseline (-60 to -50 s). n = 11 mice.

**(E)** ΔF/F activity during NREMs. The duration of NREMs episodes was normalized in time, ranging from 0 to 100%. Shading, ± s.e.m. Pairwise t-tests with Holm-Bonferroni correction p < 5.36e-33 between 20 and 100. Gray bar, intervals where ΔF/F activity was significantly different from baseline (0 to 20%, the first time bin). n = 889 events.

(**F**) Average normalized EEG spectrogram (top), and z-scored ΔF/F activity (bottom) during two successive REMs episodes and the inter-REM interval. Each REMs episode and inter-REM interval was compressed to unit length and the ΔF/F activity was averaged over multiple episodes/intervals. Shading, s.e.m.

### Figure S3

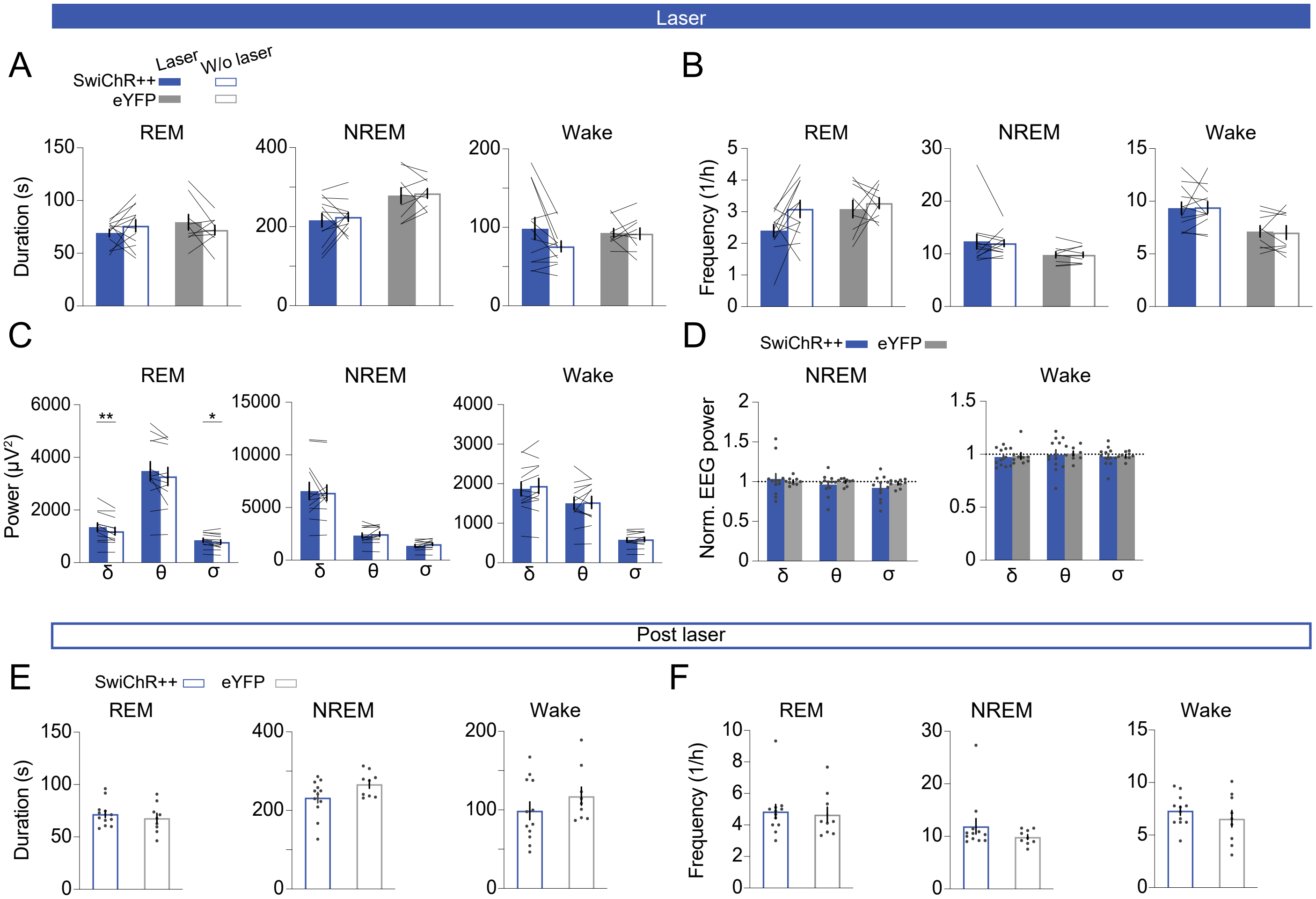

Laser
A
SwiChR++
eYFP
Laser
W/o laser
B
REM
NREM
Wake
Duration (s)
Frequency (1/h)
C
D
SwiChR++
eYFP
Power (μV²)
Norm. EEG power
δ
θ
σ
**
*
Post laser
E
SwiChR++
eYFP
F

**FIGURE S3 | Effects of inhibiting POA$^{GAD2}$→TMN neurons on brain states and EEG. Related to FIgure 2.**

**(A)** Duration of REMs, NREMs, and wake episodes with and without laser stimulation in SwiChR++ and eYFP mice.

**(B)** Frequency of REMs, NREMs, and wake episodes with and without laser stimulation in SwiChR++ and eYFP mice.

**(C)** Comparison of EEG δ, θ, and σ power during REMs, NREMs, and wakefulness with and without laser stimulation in SwiChR++ mice. Paired t-tests, SwiChR-laser vs. SwiChR-w/o laser $p = 0.0048$, 0.0158 for δ and σ during REM.

**(D)** Normalized EEG δ, θ, and σ power during NREMs, and wakefulness in SwiChR++ and eYFP mice.

**(E)** Duration of REMs, NREMs, and wake episodes during 3 h recordings directly following the laser stimulation interval (post laser) in SwiChR++ and eYFP mice.

**(F)** Frequency of REMs, NREMs, and wake episodes during post laser recordings in SwiChR++ and eYFP mice.

**(A-C)** Bars, averages across mice; lines, individual mice; error bars, ± s.e.m.

**(D-F)** Bars, averages across mice; dots, individual mice, error bars, ± s.e.m.

SwiChR++: n = 12 mice; eYFP: n = 9 mice

### Figure S4

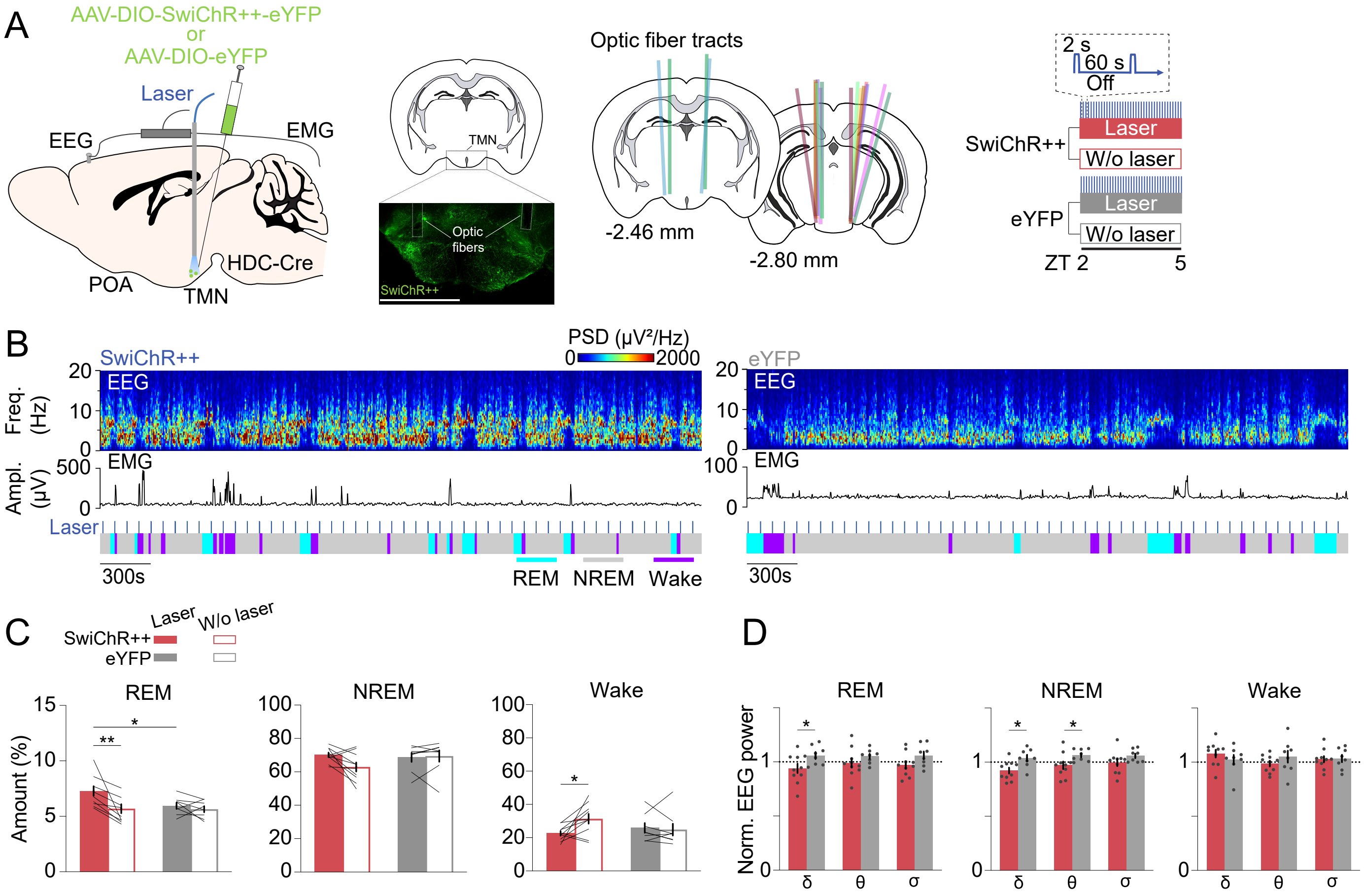

A
AAV-DIO-SwiChR++-eYFP
or
AAV-DIO-eYFP
Laser
EEG
EMG
HDC-Cre
POA
TMN
Optic fibers
SwiChR++
Optic fiber tracts
-2.46 mm
-2.80 mm
2 s
60 s
Off
SwiChR++
Laser
W/o laser
eYFP
Laser
W/o laser
ZT 2
5
B
SwiChR++
PSD (μV²/Hz)
0
2000
Freq. (Hz)
EEG
Ampl. (μV)
EMG
Laser
300s
REM
NREM
Wake
eYFP
C
Laser
W/o laser
SwiChR++
eYFP
REM
NREM
Wake
Amount (%)
D
REM
NREM
Wake
Norm. EEG power
δ
θ
σ

**FIGURE S4 | Effects of inhibiting TMN$^{HIS}$ neurons on brain states and EEG. Related to Figure 2.**

(**A**) Left, schematic of optogenetic inhibition experiments. Center left, fluorescence image of TMN in HDC-Cre mouse injected with $AAV_2$-EF1a-DIO-SwiChR++-eYFP into the TMN. Scale bar, 1 mm. Center right, location of optic fiber tracts. Each colored bar represents the location of an optic fiber. Right, experimental paradigm for laser stimulation (2 s step pulses at 60s intervals) in SwiChR++ and eYFP-expressing mice.
(**B**) Example recording of a SwiChR++ (left) and eYFP mouse (right) with laser stimulation. Shown are EEG power spectra, EMG amplitude, and color-coded brain states.
(**C**) Percentage of time spent in REMs, NREMs, and wakefulness with and without laser in SwiChR++ and eYFP mice. REMs: Mixed ANOVA, virus $p = 0.1470$, laser $p = 0.0054$, interaction $p = 0.0455$; t-tests with Bonferroni correction, SwiChR-laser vs. SwiChR-w/o laser $p = 0.0019$, SwiChR-laser vs. eYFP-laser $p = 0.0384$. Wake: Mixed ANOVA, virus $p = 0.6180$, laser $p = 0.0909$, interaction $p = 0.0378$; t-tests with Bonferroni correction, SwiChR-laser vs. SwiChR-w/o laser $p = 0.0154$.
**(D**) Normalized EEG δ, θ, and σ power during REMs, NREMs, and wake in SwiChR++ and eYFP mice with laser. Unpaired t-tests, REMs: SwiChR vs. eYFP $p = 0.0498$ for δ; NREMs: SwiChR vs. eYFP $p = 0.0123$, $0.0474$ for δ and θ.

(**C**) Bars, averages across mice; lines, individual mice; error bars, ± s.e.m.
(**D**) Bars, averages across mice; dots, individual mice; error bars, ± s.e.m.

SwiChR++: n = 10 mice; eYFP: n = 8 mice

### Figure S5

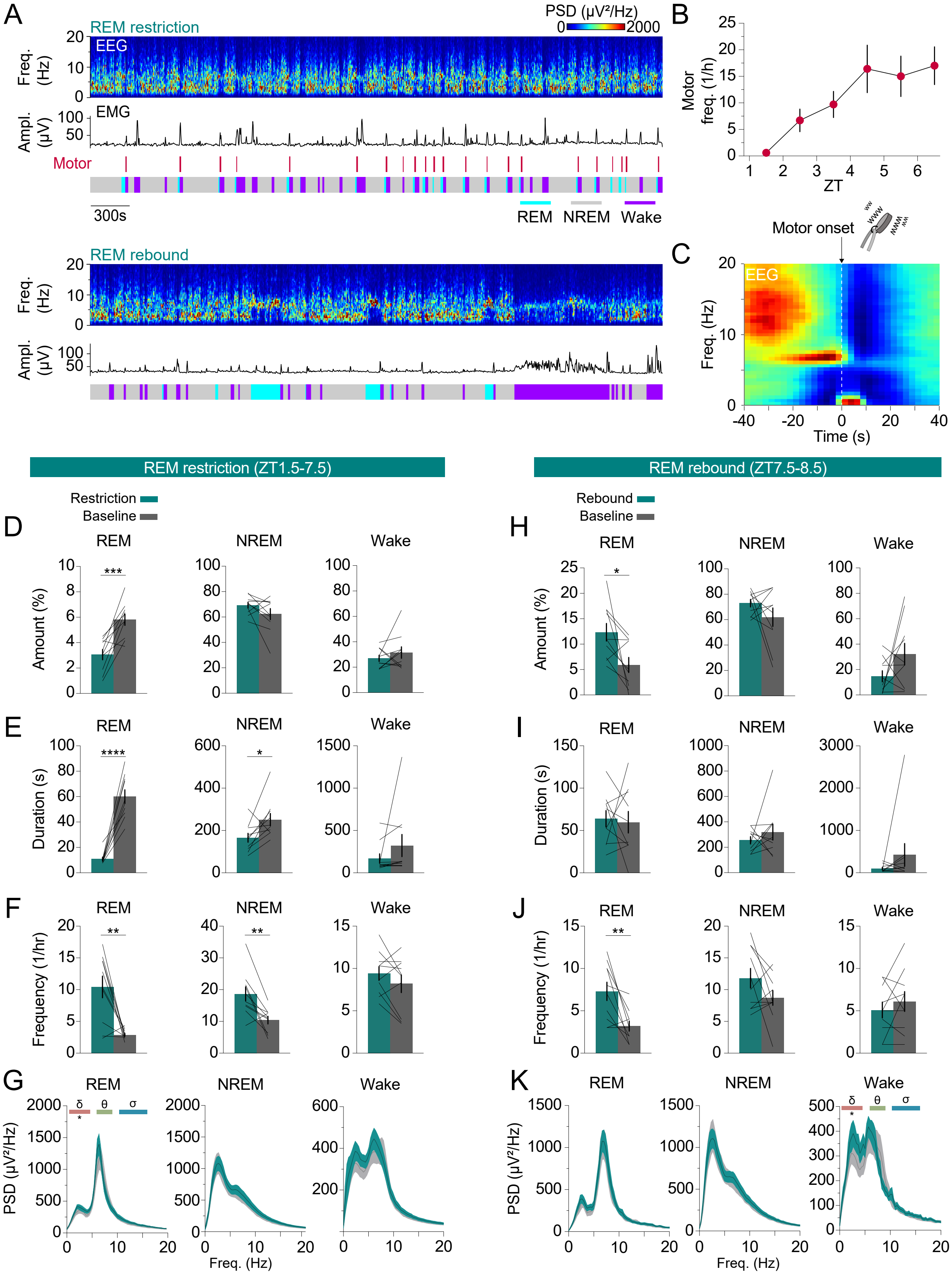
A
REM restriction
PSD (µV²/Hz)
0
2000
EEG
Freq. (Hz)
Ampl. (µV)
EMG
Motor
300s
REM
NREM
Wake
REM rebound
B
Motor freq. (1/h)
ZT
C
Motor onset
EEG
Freq. (Hz)
Time (s)
REM restriction (ZT1.5-7.5)
REM rebound (ZT7.5-8.5)
Restriction
Baseline
Rebound
D
E
F
G
H
I
J
K
Amount (%)
Duration (s)
Frequency (1/hr)
PSD (µV²/Hz)
Freq. (Hz)
δ
θ
σ

**FIGURE S5 | Effects of REMs restriction on brain states and EEG. Related to Figure 3.**

**(A)** Example session during REMs restriction (top) and REMs rebound (bottom) from the same mouse. Shown are EEG spectrogram, EMG amplitude, motor vibration events, and color-coded brain states.
**(B)** Frequency of motor vibration events throughout REMs restriction (ZT 1.5-7.5). Error bars, ± s.e.m. One-way ANOVA $p = 0.0061$.
**(C)** Average normalized EEG spectrogram preceding motor vibration onset.
**(D)** Percentage of time spent in REMs, NREMs, and wakefulness during 6 h of REMs restriction (green, ZT 1.5-7.5) and baseline recordings (gray, ZT 1.5-7.5). Paired t-test, $p = 0.0006$ for REMs amount.
**(E)** Duration of REMs, NREMs, and wake episodes during 6 h of REMs restriction (green) and baseline recordings (gray) (ZT 1.5-7.5). Paired t-tests, p = 1.44e-5, 0.0276 for the duration of REMs and NREMs episodes.
**(F)** Frequency of REMs, NREMs, and wake episodes during 6 h of REMs restriction (green) and baseline recordings (gray)(ZT 1.5-7.5). Paired t-tests, $p = 0.0027$ and 0.0043 for the frequency of REMs and NREMs episodes.
**(G)** Power spectral density (PSD) of EEG during REMs, NREMs, and wakefulness during 6 h of REMs restriction (green) and baseline recordings (gray)(ZT 1.5-7.5). Paired t-tests, $p = 0.0293$ for REMs $\delta$ power.
**(H)** Percentage of time spent in REMs, NREMs, and wakefulness during 1 h of REMs rebound (green, ZT 7.5-8.5) and baseline sleep (gray, ZT 7.5-8.5). Paired t-tests, $p = 0.0226$ for REMs amount.
**(I)** Duration of REMs, NREMs, and wake episodes during 1 h of REMs rebound (green) and baseline recordings (gray) (ZT 7.5-8.5).
**(J)** Frequency of REMs, NREMs, and wake episodes during 1 h of REMs rebound (green) and baseline recordings (gray). Paired t-tests, $p = 0.0055$ for the frequency of REMs episodes (ZT 7.5-8.5).
**(K)** PSD of EEG during REMs, NREMs, and wakefulness during 1 h of REMs rebound (green) and baseline recordings (gray). Paired t-tests, $p = 0.0164$ for wake $\delta$ power (ZT 7.5-8.5).

**(D-F, H-J)** Bars, averages across mice; lines, individual mice; error bars, ± s.e.m. n = 10 mice
**(G, K)** Shadings, ± s.e.m. n = 10 mice in G; n = 9 mice in K, 1 mouse was excluded for REMs because there was no REMs during baseline recordings

### Figure S6

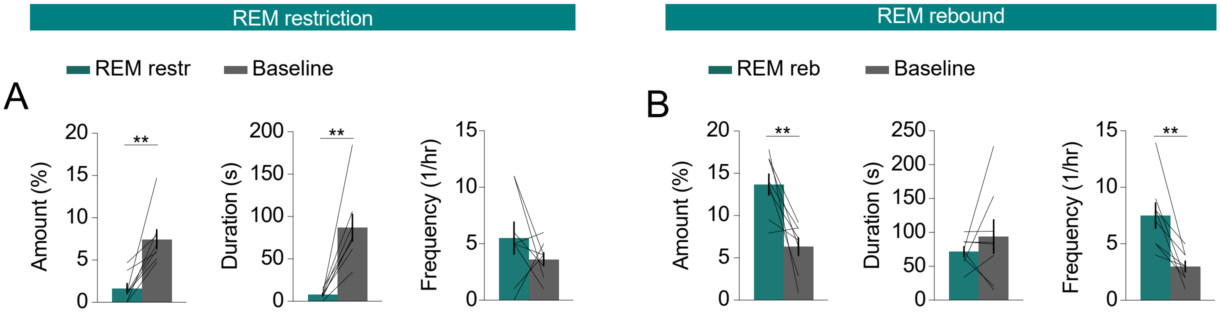

REM restriction
REM rebound
REM restr
Baseline
REM reb
Baseline
A
B
Amount (%)
Duration (s)
Frequency (1/hr)
**

**FIGURE S6 | REMs amount, duration, and frequency of episodes during photomery recordings combined with REMs restriction and rebound. Related to Figure 3.**

**(A)** Percentage of REMs, duration, and frequency of REMs episodes during REMs restriction (green, ZT 6.5-7.5) and baseline recordings (gray, ZT 6.5-7.5). Paired t-tests, $p = 0.0024, 0.0013$ for amount, and duration.

**(B)** Percentage of REMs, duration, and frequency of REMs episodes during REMs rebound (green, ZT 7.5-8.5) and baseline recordings (gray, ZT 7.5-8.5). Paired t-tests, $p = 0.0066, 0.0047$ for amount and frequency.

Bars, averages across mice; lines, individual mice; error bars, ± s.e.m.

n = 8 mice

### Figure S7

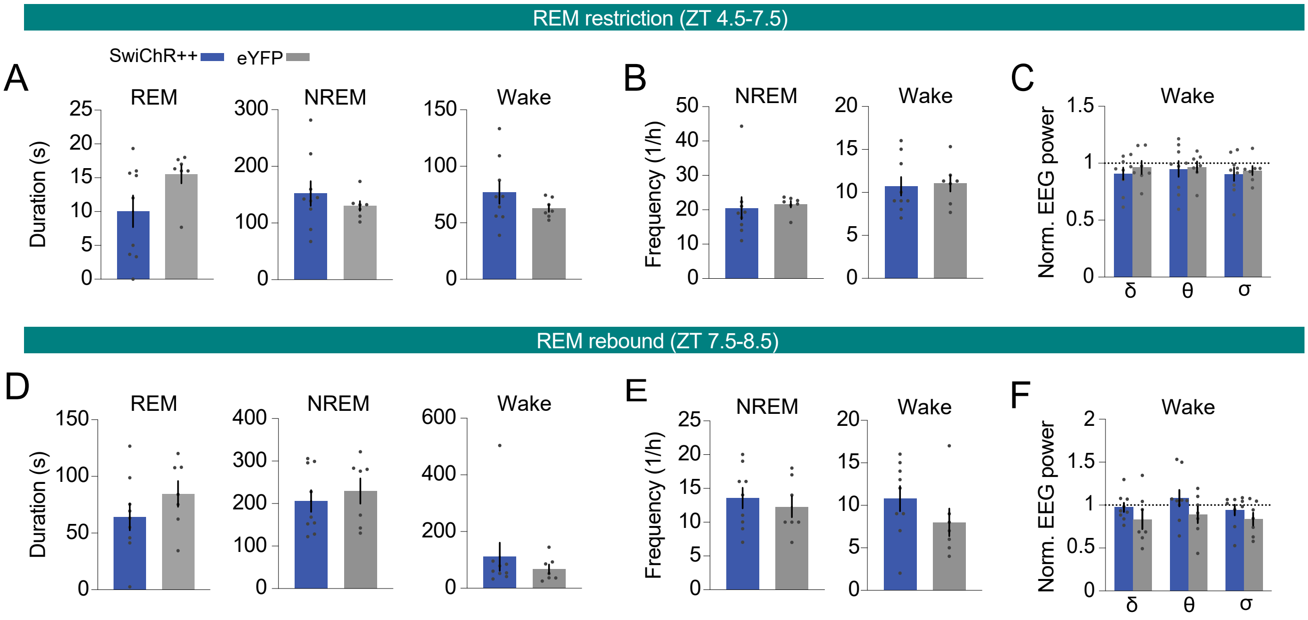
REM restriction (ZT 4.5-7.5)
SwiChR++
eYFP
A
B
C
REM
NREM
Wake
Duration (s)
Frequency (1/h)
Norm. EEG power
δ
θ
σ
REM rebound (ZT 7.5-8.5)
D
E
F

**FIGURE S7 | Duration and frequency of REMs episodes and EEG power during SwiChR++-mediated inhibition combined with REMs restriction and rebound. Related to Figure 4.**

**(A)** Duration of REMs, NREMs, and wake episodes during the last 3 h of REMs restriction with laser stimulation (ZT 4.5-7.5) in mice expressing SwiChR++ and eYFP.
**(B)** Frequency of REMs, NREMs, and wake episodes during the last 3 h of REMs restriction with laser stimulation in mice expressing SwiChR++ and eYFP.
**(C)** Normalized EEG δ, θ, and σ power during the last 3 h of REMs restriction with laser stimulation in eYFP and SwiChR++ mice.
**(D)** Duration of REMs, NREMs, and wake episodes during REMs rebound (ZT 7.5-8.5) in mice expressing SwiChR++ and eYFP.
**(E)** Frequency of REMs, NREMs, and wake episodes during REMs rebound (ZT 7.5-8.5) in mice expressing SwiChR++ and eYFP.
**(F)** Normalized EEG δ, θ, and σ power during REMs rebound in eYFP and SwiChR++ mice.

Bars, averages across mice; dots, individual mice; error bars, ± s.e.m.
SwiChR++: n = 9 mice; eYFP: n = 7 mice
